# Analysis of key genes and pathways associated with the pathogenesis of Type 2 diabetes mellitus

**DOI:** 10.1101/2021.08.12.456106

**Authors:** Varun Alur, Varshita Raju, Basavaraj Vastrad, Chanabasayya Vastrad, Shivakumar Kotturshetti

## Abstract

Type 2 diabetes mellitus (T2DM) is the most common endocrine disorder which poses a serious threat to human health. This investigation aimed to screen the candidate genes differentially expressed in T2DM by bioinformatics analysis. The expression profiling by high throughput sequencing of GSE81608 dataset was retrieved from the gene expression omnibus (GEO) database and analyzed to identify the differentially expressed genes (DEGs) between T2DM and normal controls. Then, Gene Ontology (GO) and pathway enrichment analysis, protein-protein interaction (PPI) network, modules, miRNA-hub gene regulatory network construction and TF-hub gene regulatory network construction, and topological analysis were performed. Receiver operating characteristic curve (ROC) analysis was also performed to verify the diagnostics value and expression of identified hub genes. A total of 927 DEGs (461 were up regulated and 466 down regulated genes) were identified in T2DM. GO and REACTOME results showed that DEGs mainly enriched in protein metabolic process, establishment of localization, metabolism of proteins and metabolism. The top centrality hub genes APP, MYH9, TCTN2, USP7, SYNPO, GRB2, HSP90AB1, UBC, HSPA5 and SQSTM1 were screened out as the critical genes among the DEGs from the PPI network, modules, miRNA-hub gene regulatory network construction and TF-hub gene regulatory network. ROC analysis provide diagnostics value of hub genes. This study identified key genes, signal pathways and therapeutic agents, which might help us, improve our understanding of the mechanisms of HGPS and identify some new therapeutic agents for T2DM.

## Introduction

Type 2 diabetes mellitus (T2DM) is a complex metabolic disorder and is characterized primarily by a decrease in insulin secretion, typically accompanied by insulin resistance [1]. Globally, it is predicted that 25 million adults (20–79 years) have diabetes, projected to reach 629 million by 2045 and is the ninth leading cause of death [2-3]. T2DM is mainly associated with macrovascular complications include stroke, coronary artery disease and peripheral arterial disease, and microvascular include diabetic retinopathy, diabetic nephropathy, and diabetic neuropathy, and non-vascular diabetes complications include nonalcoholic fatty liver disease, psychiatric disease, obesity, cancer, cognitive impairment, infections and disability [4]. There are several important risk factors for T2DM, such as age, sex, family history of diabetes, hypertension, obesity, abdominal obesity, stress in the workplace or home, a sedentary lifestyle, smoking, insufficient fruit and vegetable consumption, physical activity, genetic and environmental [5]. Our understanding of the occurrence and development mechanism of T2DM has been greatly improved; however, the cause and potential molecular mechanism of T2DM are still unclear [6]. Therefore, it is necessary to identify key genes and pathways for understanding the molecular mechanism and discovering potential biomarkers for T2DM.

In recent decades, more and more researchers have devoted themselves to exploring the potential mechanisms for progression of T2DM. For instance, it has been demonstrated that the HHEX, CDKN2A/B, and IGF2BP2 [7], CDKAL1 and HHEX/IDE [8], ADIPOQ, PPAR-γ and RXR-α [9], ABCC8 and KCNJ11 [10], TCF7L2, SLC30A8, PCSK1 and PCSK2 [11], PI3K/AKT-and AMPK signaling pathway [12], mTOR signaling pathway [13], insulin signaling pathway [14], AGE/RAGE/JNK, STAT3/SCOS3 and RAS signaling pathway [15] and ERK signaling pathway [16] were involved in progression of T2DM. Therefore, it is of great practical significance to explore the genes and signaling pathways of T2DM on islet cells.

RNA sequencing technology can rapidly detect gene expression on a global basis and are particularly useful in screening for differentially expressed genes (DEGs) in diseases [17]. RNA sequencing which allow the investigation of gene expression in a high throughput manner with high sensitivity, specificity and repeatability. Significant amounts of data have been produced via the use of RNA sequencing and the majority of such data has been uploaded and stored in public databases. Previous investigations concerning T2DM gene expression profiling have identified hundreds of DEGs [18]. However, the comparative analysis of DEGs across a range of independent investigation might yield only a relatively limited amount of useful data with regard to T2DM advancement. The disadvantages of these single investigations might be overcome by bioinformatics analysis, as this approach would make it possible to analyze the signaling pathways and interaction networks linked with the identified DEGs. This knowledge might help in elucidating the molecular mechanisms underlying T2DM.

In the present investigation, expression profiling by high throughput sequencing dataset was downloaded from the Gene Expression Omnibus (GEO) (GEO, http://www.ncbi.nlm.nih.gov/geo/) [19]: GSE81608 [20]. DEGs were identified in T2DM. Additionally, gene ontology (GO), REACTOME pathway enrichment analysis was performed and protein-protein interaction (PPI) networks, modules, miRNA-hub gene regulatory network construction and TF-hub gene regulatory network were constructed to identify the hub genes, miRNA and TFs in T2DM. Finally, hub genes were validated by receiver operating characteristic curve (ROC). Collectively, the findings of the present investigation highlighted crucial genes and signaling pathways that might contribute to the pathology of T2DM. These may provide a basis for the advancement of future diagnostic, prognostic and therapeutic tools for T2DM.

## Materials and Methods

### Data resources

Expression profiling by high throughput sequencing dataset GSE81608 [20] was downloaded from the GEO database. The data was produced using a GPL16791 Illumina HiSeq 2500 (Homo sapiens). The GSE81608 dataset contained data from 1600 samples, including 949 T2DM, and 651 healthy control samples.

### Identification of DEGs

Limma package in R software [21] is a tool to identify DEGs by comparing samples from GEO series. Limma package in R software was used to search for in messenger RNAs (mRNAs; DEGs) that were differentially expressed between T2DM and healthy control samples. The cutoff criteria were an adjusted p-value of <0.05, whereas the logFC value were > 0.181 for up regulated genes and < - 0.27 for down regulated genes. DEG of this dataset was visualized with volcano map and hierarchical clustering heat map. The volcano plot was drawn using ggplot2 package in R software. Hierarchical clustering heat maps of DEG expression (up regulated genes and down regulated genes) were visualized with gplots package in R software.

### GO and REACTOME pathway enrichment analysis of DEGs

GO enrichment analysis (http://geneontology.org/) [22] implement the annotation of biological processes (BP), cellular components (CC) and molecular functions (MF) of DEGs. REACTOME (https://reactome.org/) [23] is a database that stores large amounts of data on genomics, biological pathways, signaling pathways, diseases, drugs, and chemicals. The present investigation used Database for g:Profiler (http://biit.cs.ut.ee/gprofiler/) [24] to perform GO and REACTOME pathway enrichment analysis. P<0.05 was considered to indicate a statistically significant difference.

### Construction of the PPI network and module analysis

The IID interactome database (http://iid.ophid.utoronto.ca/) may be searched for associations between known and predicted proteins, and is commonly used to predict PPI information in molecular biology [25]. Cytoscape 3.8.2 (http://www.cytoscape.org/) [26] was used to visualize the results from the PPI network In this investigation, node degree [27], betweenness centrality [28], stress centrality [29] and closeness centrality [30], which constitutes a fundamental parameter in network theory, was adopted to evaluate the nodes in a network. The node degree betweenness centrality, stress centrality and closeness centrality methods were calculated using Cytoscape plugin Network Analyzer. Module analysis on the PPI network results was performed using the PEWCC1 (http://apps.cytoscape.org/apps/PEWCC1) [31] clustering algorithm that comes with Cytoscape. Module analysis might be used to find out more connected gene groups. In addition, the module analysis were further analyzed for GO and pathway enrichment analysis.

### MiRNA-hub gene regulatory network construction

Prediction of miRNA-hub genes was performed by miRNet database (https://www.mirnet.ca/) [32]. According to the regulatory interaction, miRNA-hub gene regulatory network was constructed based on miRNet by Cytoscape 3.8.2 software [26].

### TF-hub gene regulatory network construction

Prediction of TF-hub genes was performed by NetworkAnalyst database (https://www.networkanalyst.ca/) [33]. According to the regulatory interaction, TF-hub gene regulatory network was constructed based on NetworkAnalyst by Cytoscape 3.8.2 software [26].

### Validation of hub genes by receiver operating characteristic curve (ROC) analysis

ROC curve analysis was performed to evaluate the sensitivity and specificity of the hub genes for T2DM diagnosis using the R package “pROC” [34]. An area under the curve (AUC) value was determined and used to label the ROC effect.

## Results

### Identification of DEGs

A total of 927 genes were identified to be differentially expressed between T2DM and normal control samples with the threshold of adjusted p-value of <0.05, and logFC value were > 0.181 for up regulated genes and < - 0.27 for down regulated genes. Among these DEGs, 461 were up regulated and 466 down regulated genes in T2DM compared with normal control samples and are listed in Table 1. A heat map (Fig. 1) and a volcano plot (Fig. 2) for the identified DEGs was generated.

**Table 1.**
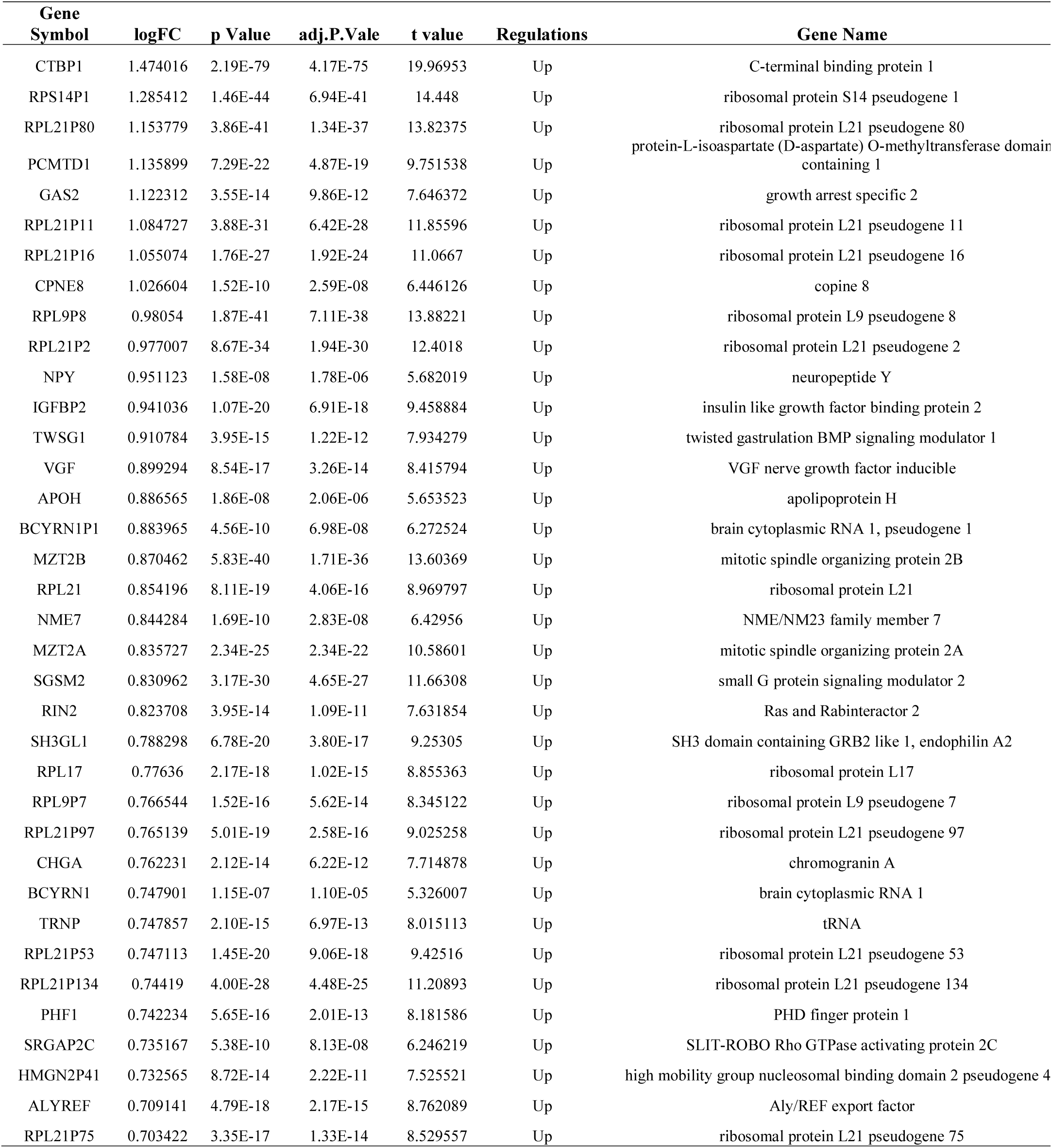
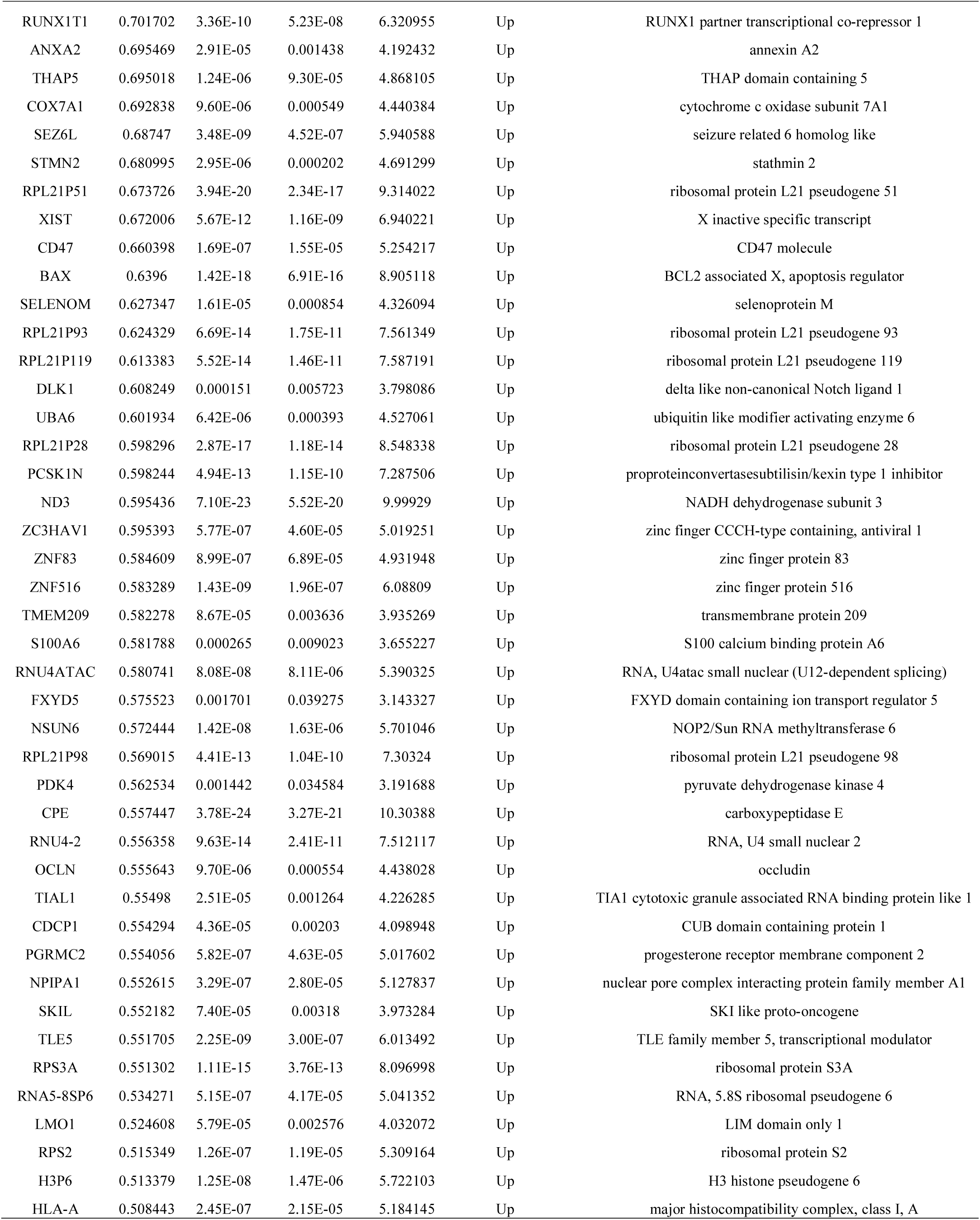
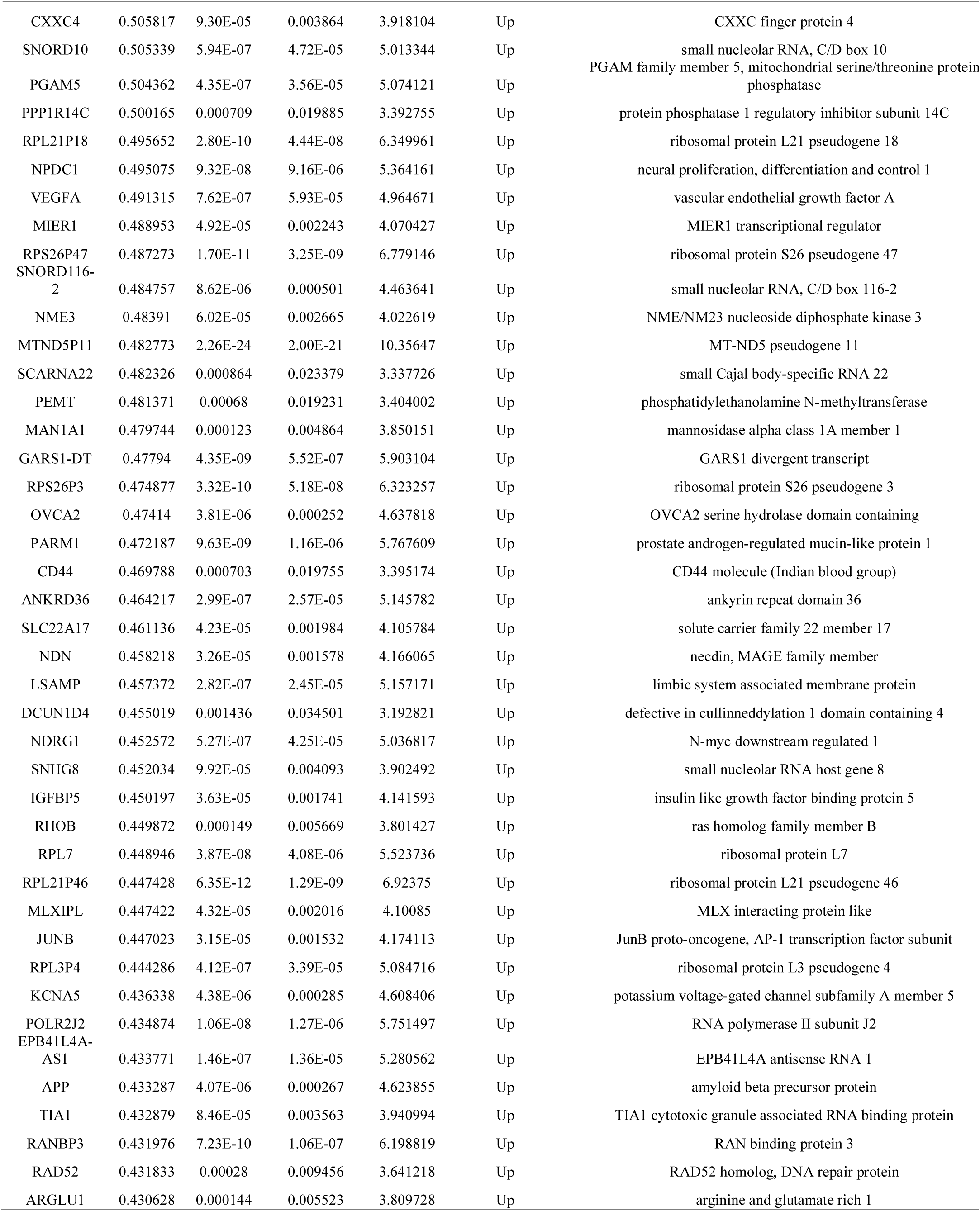
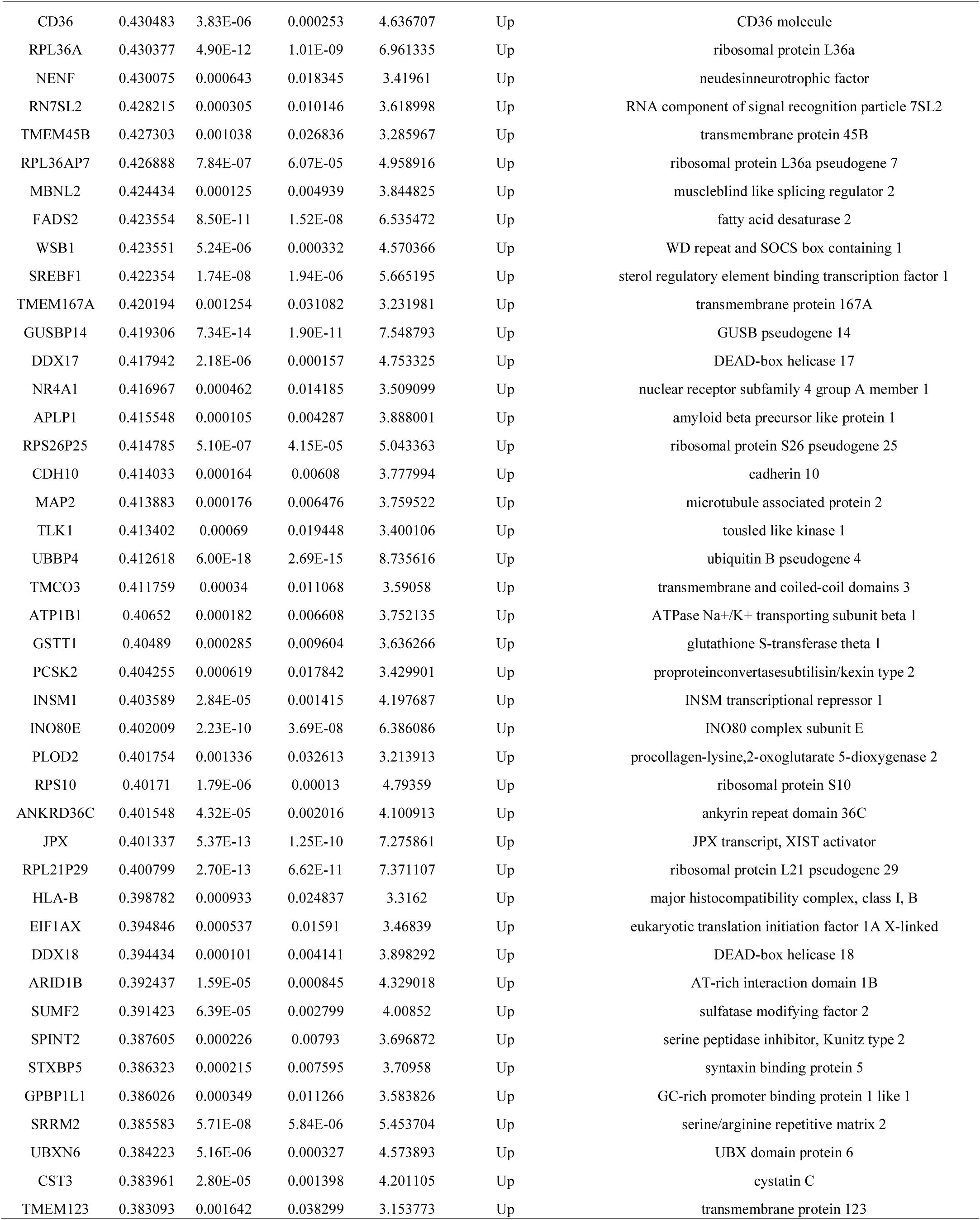
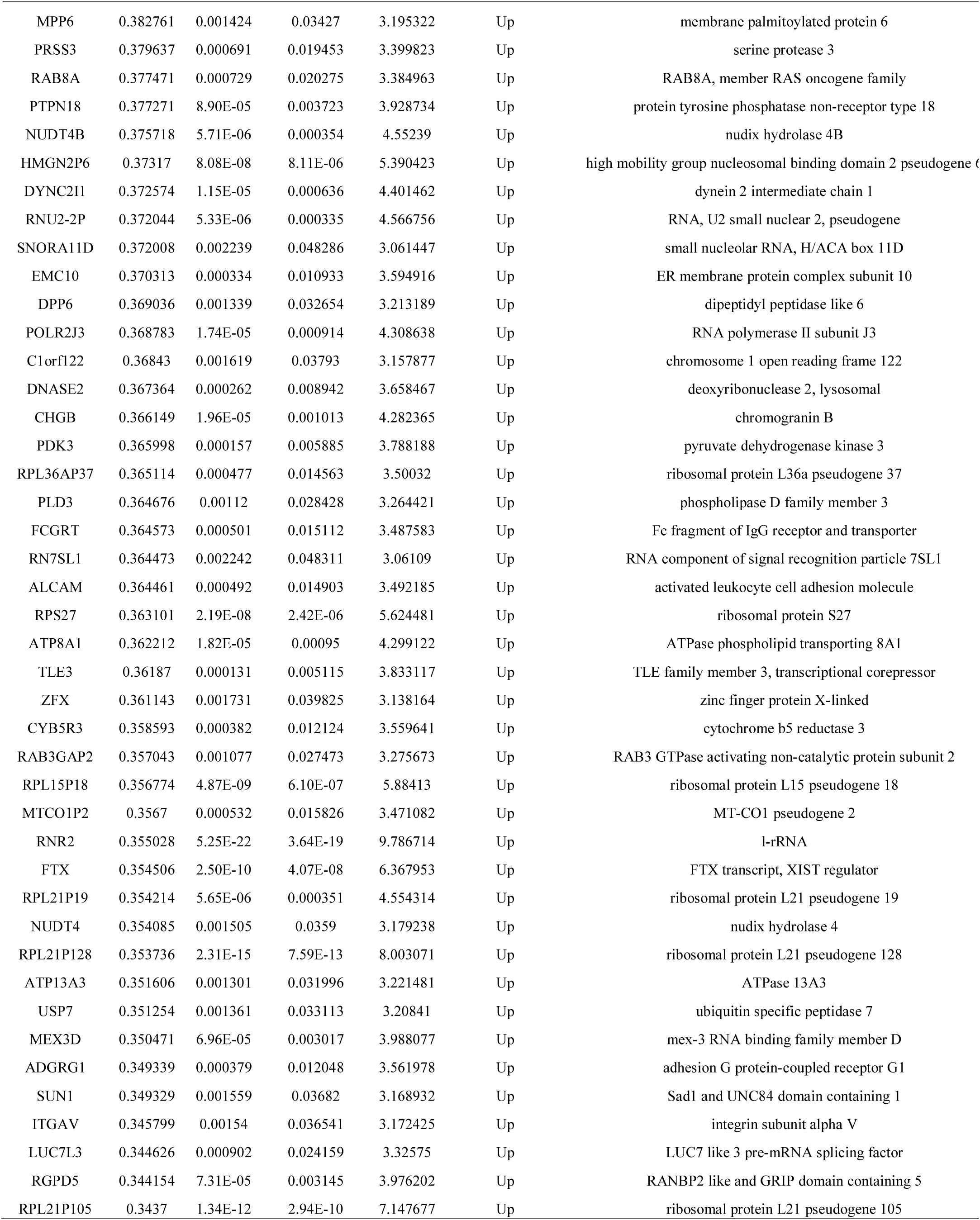
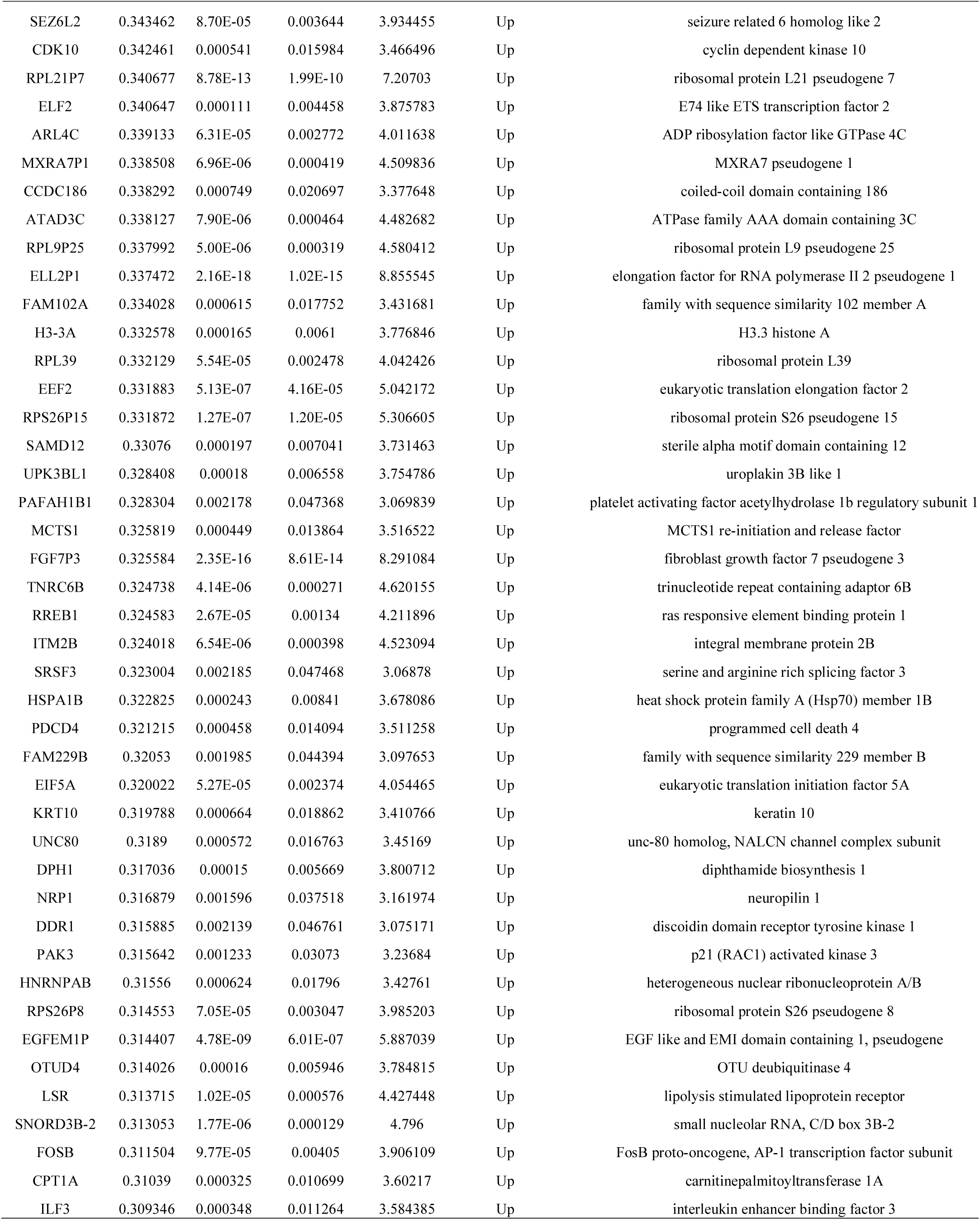
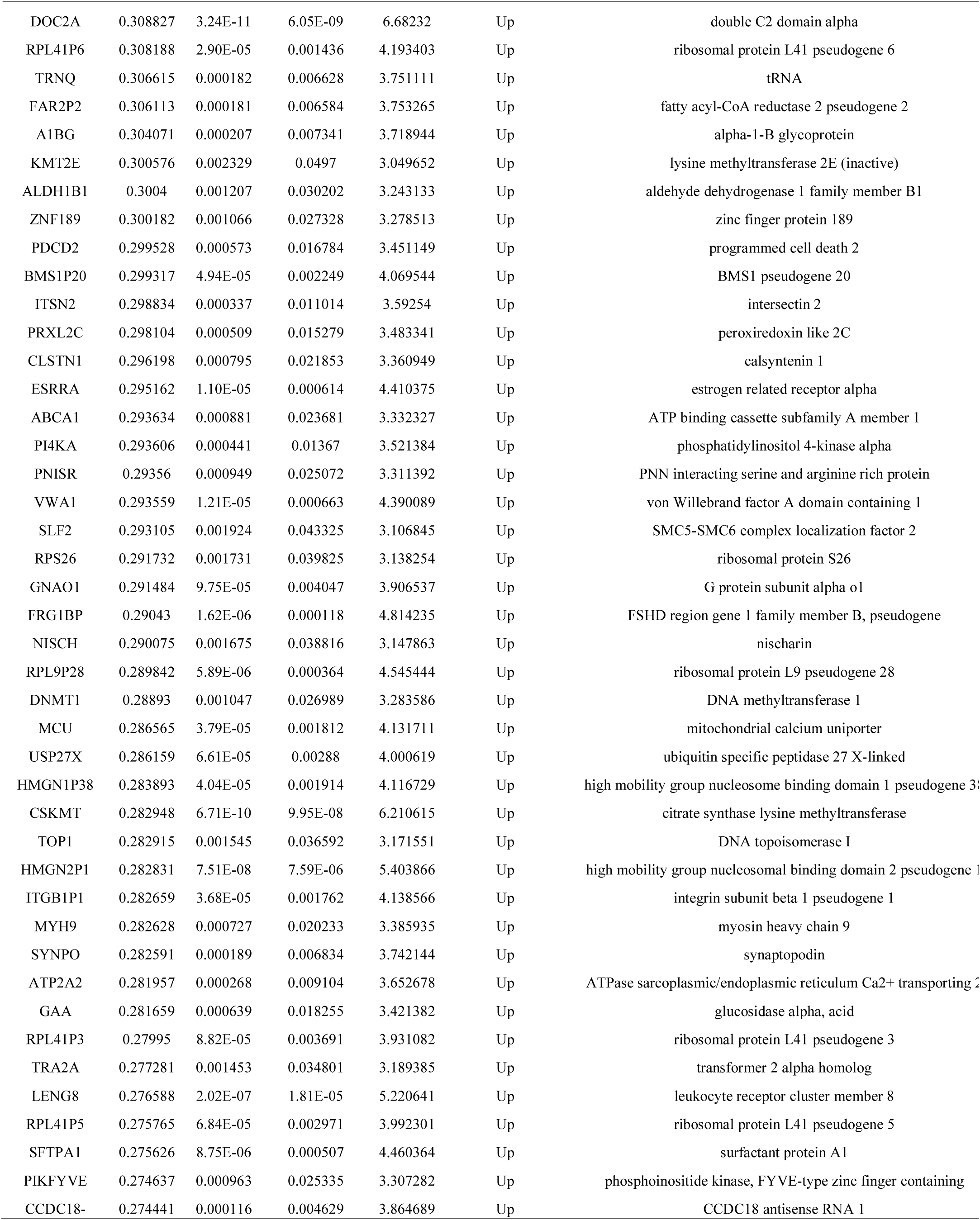
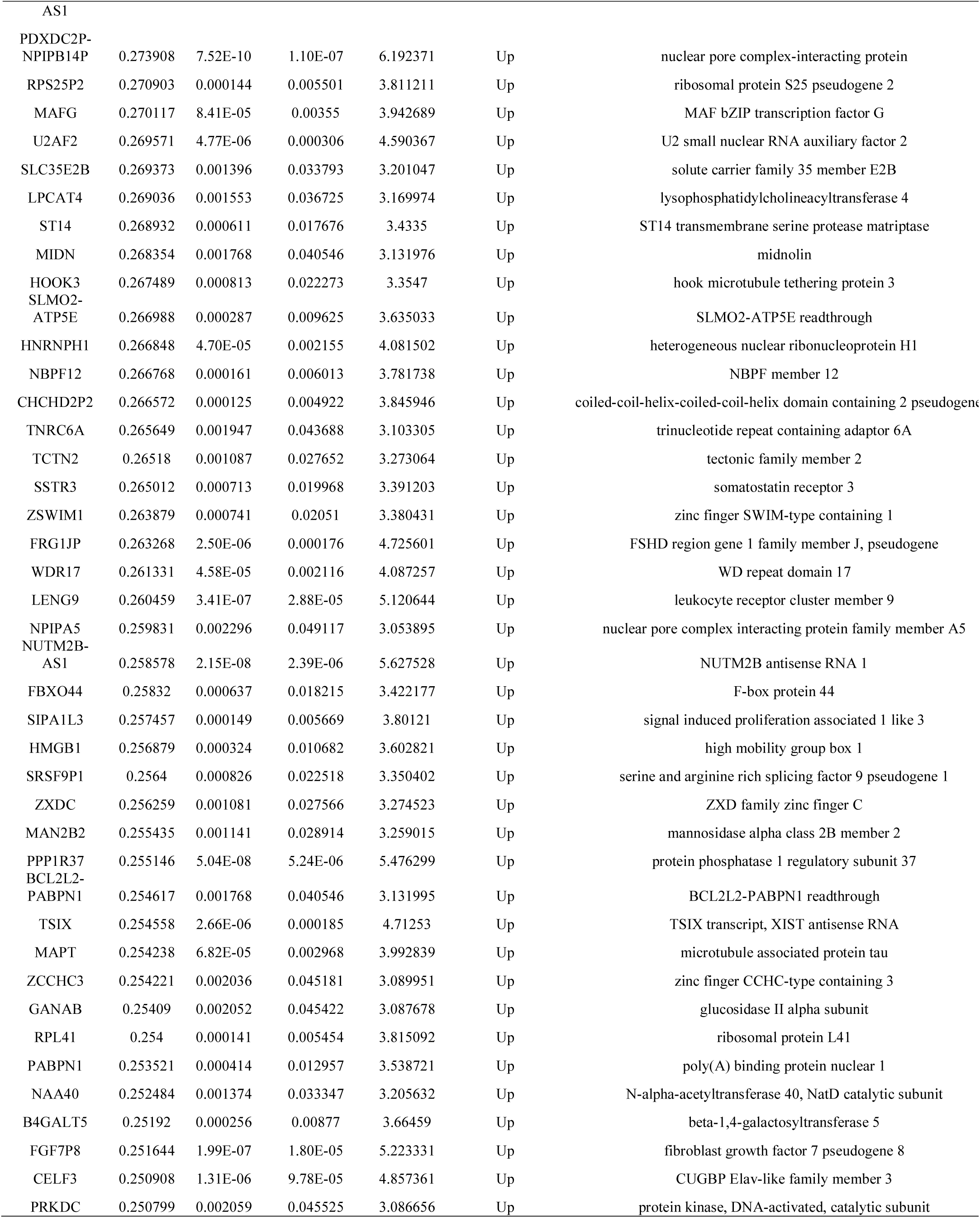
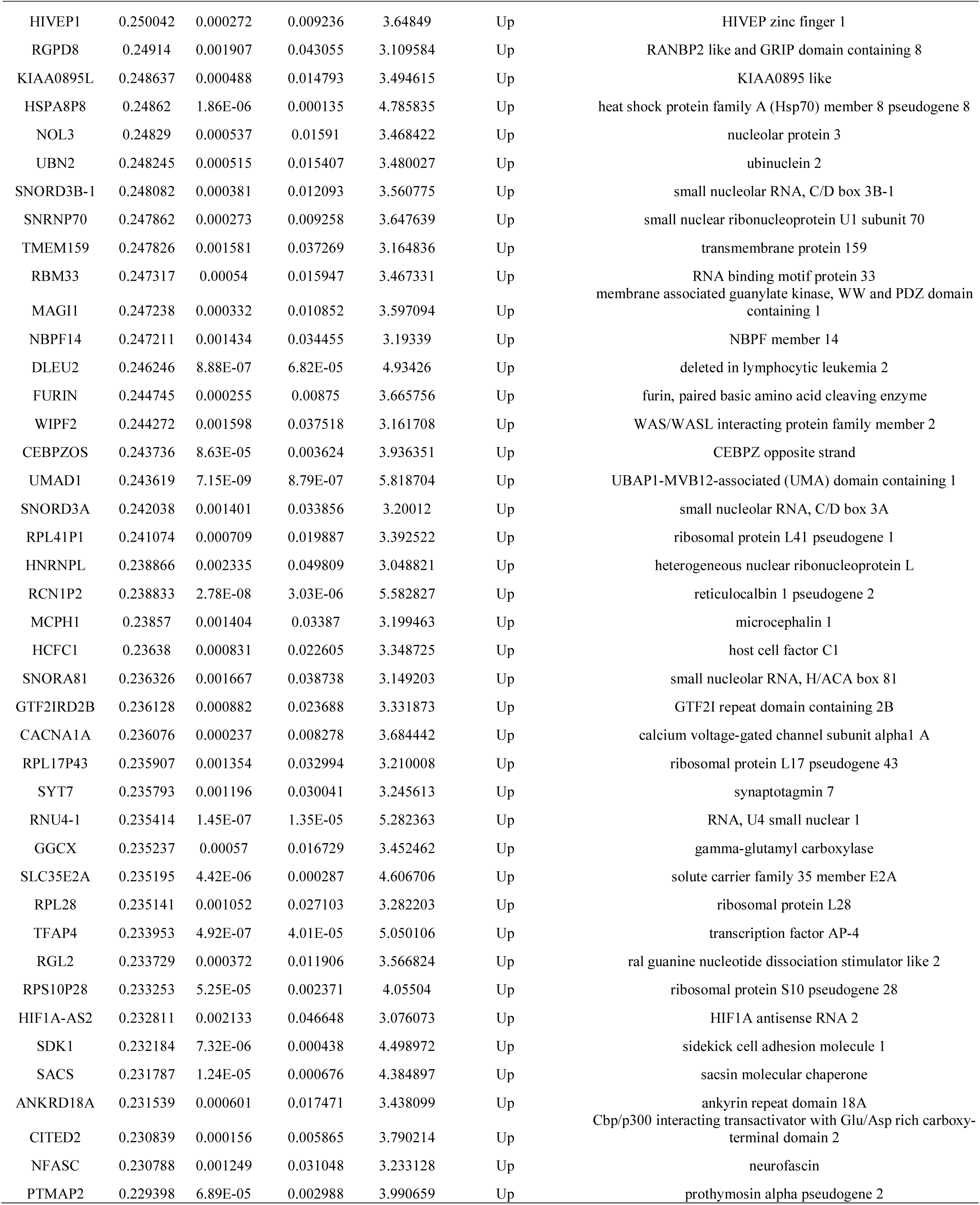
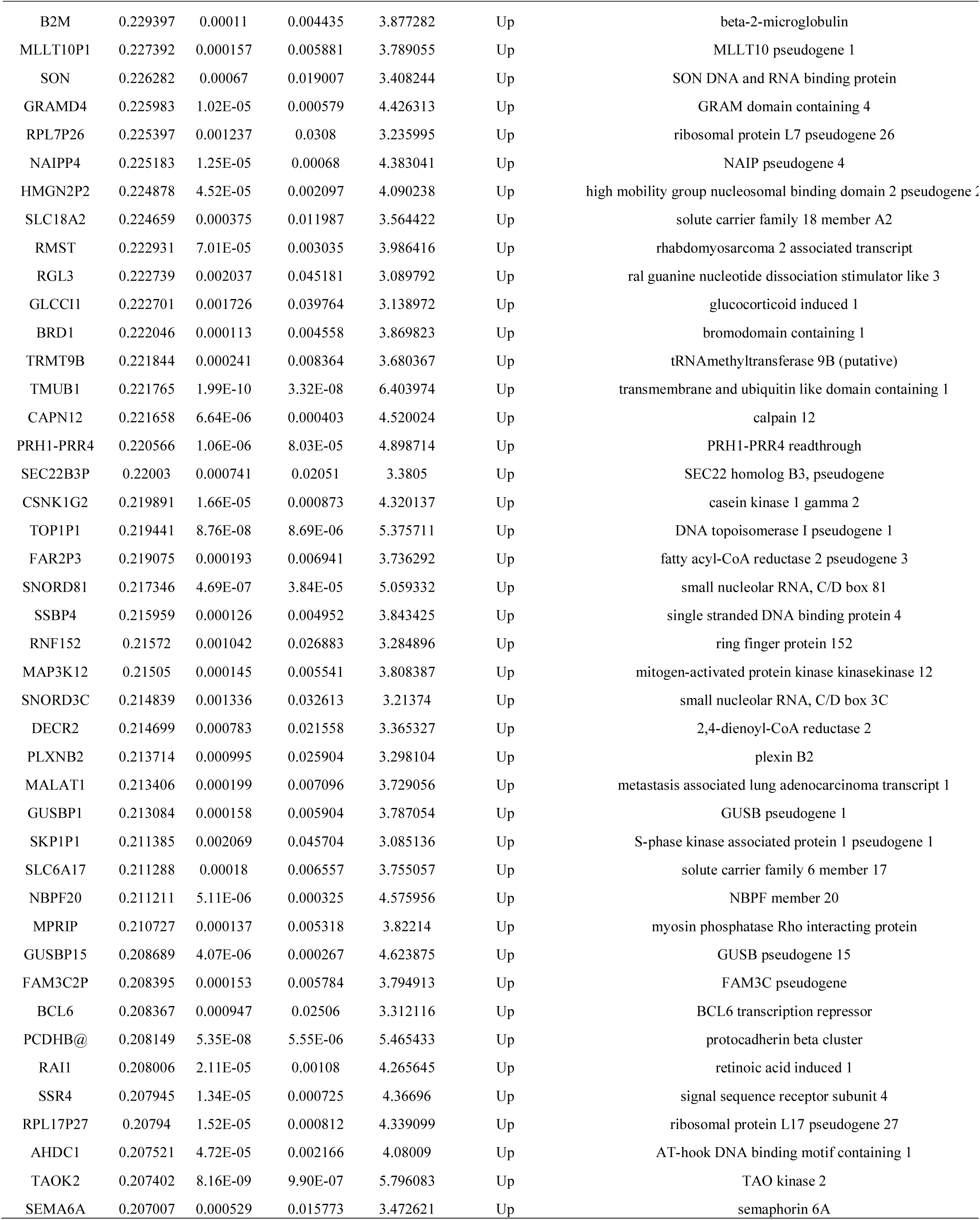
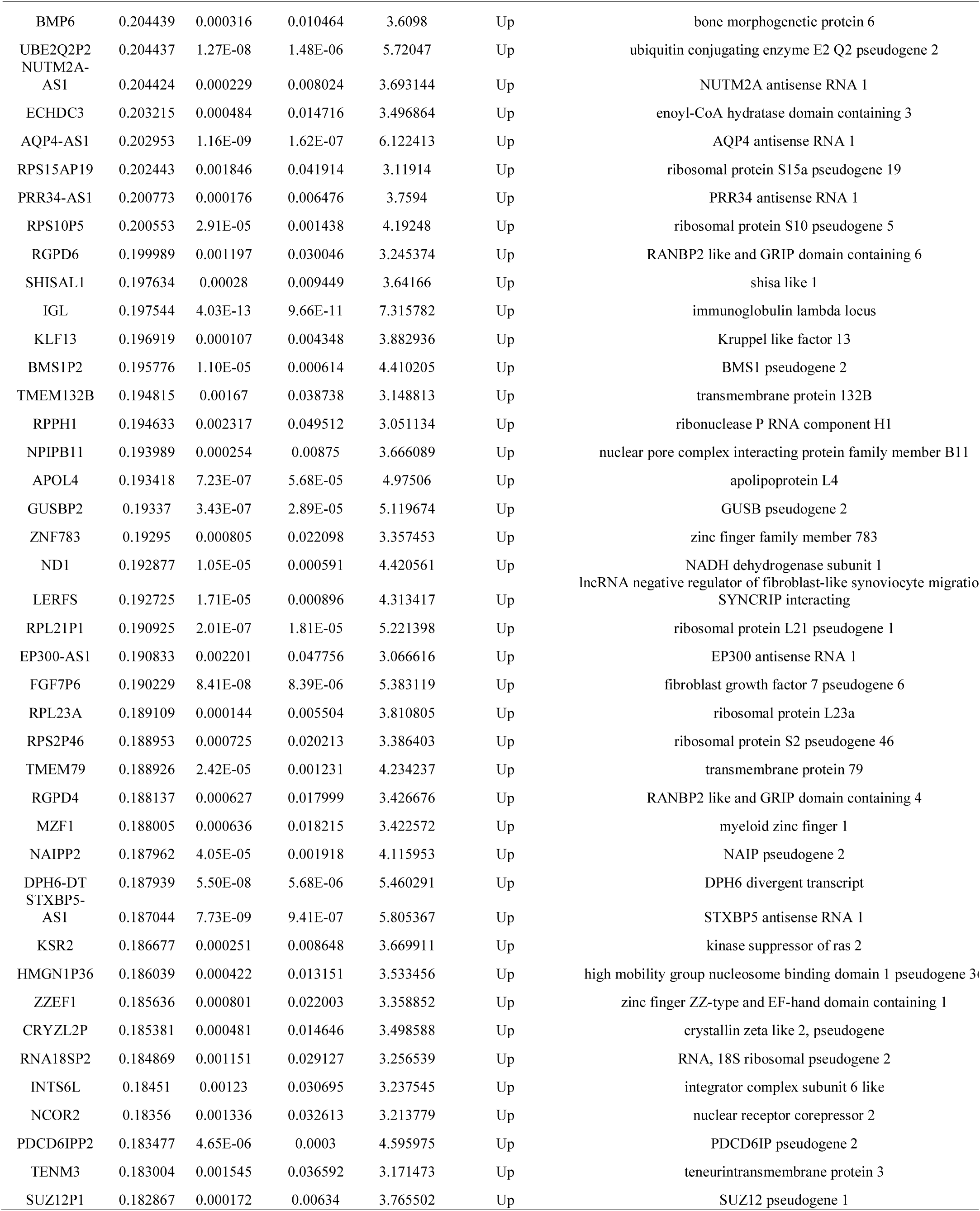
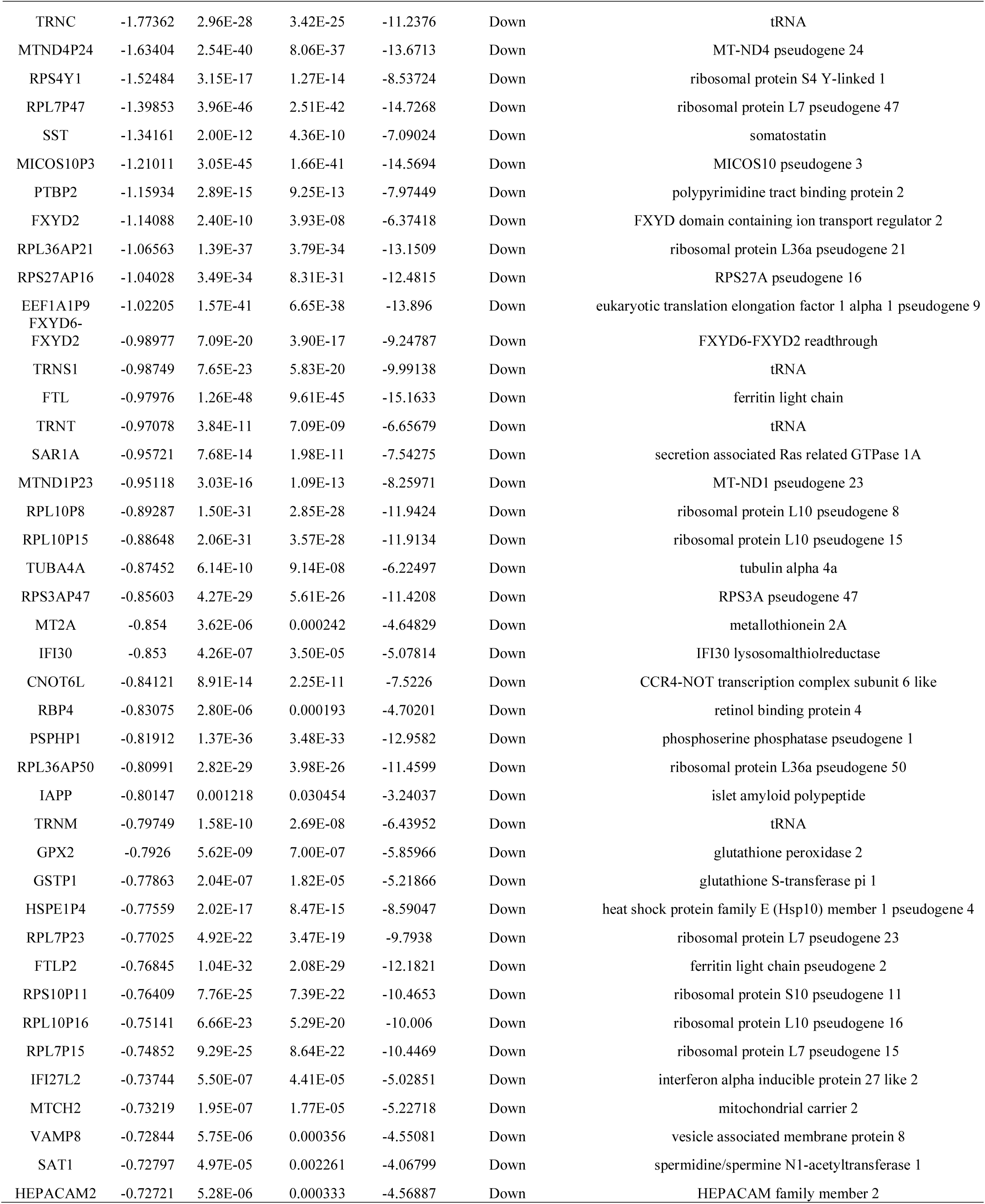
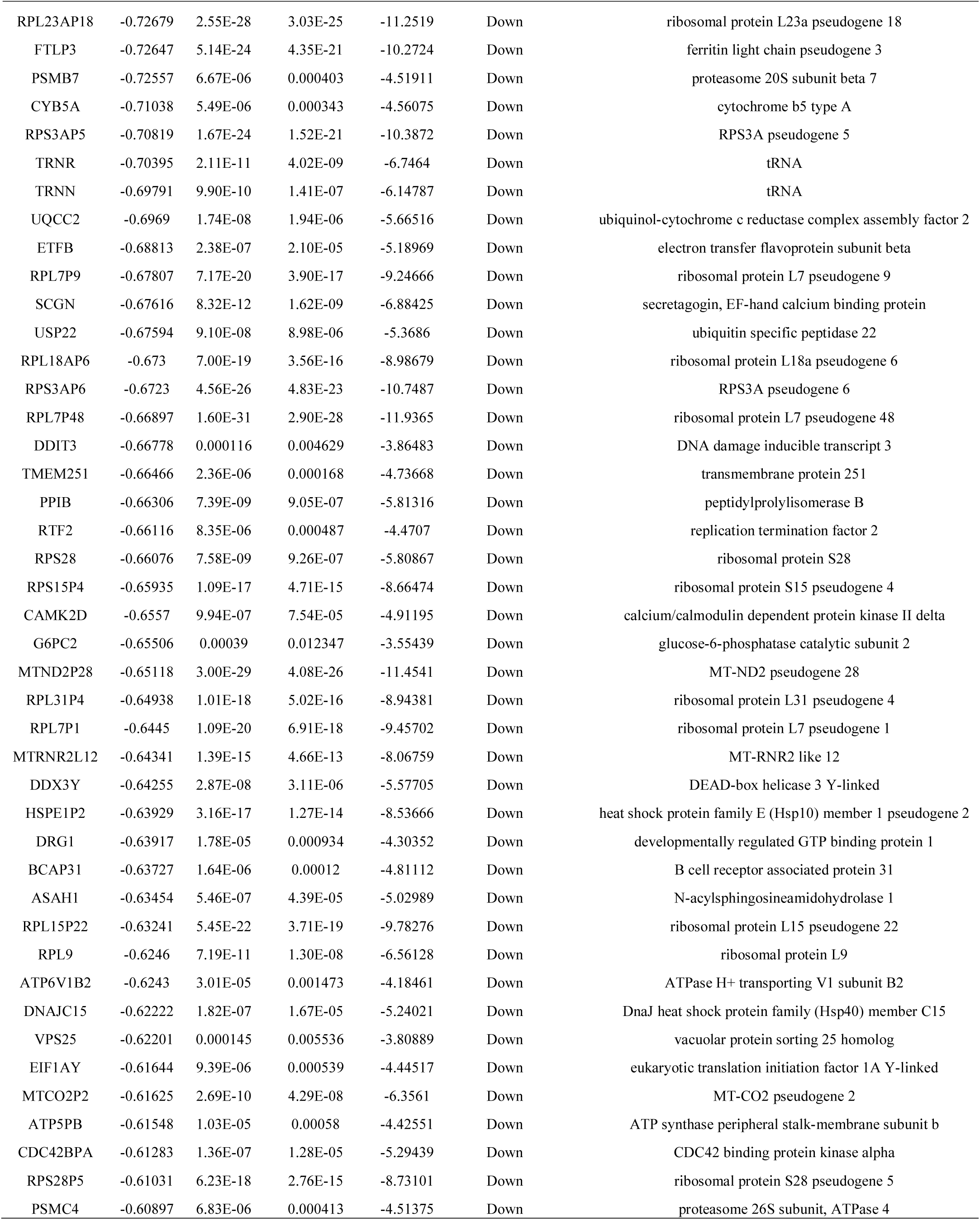
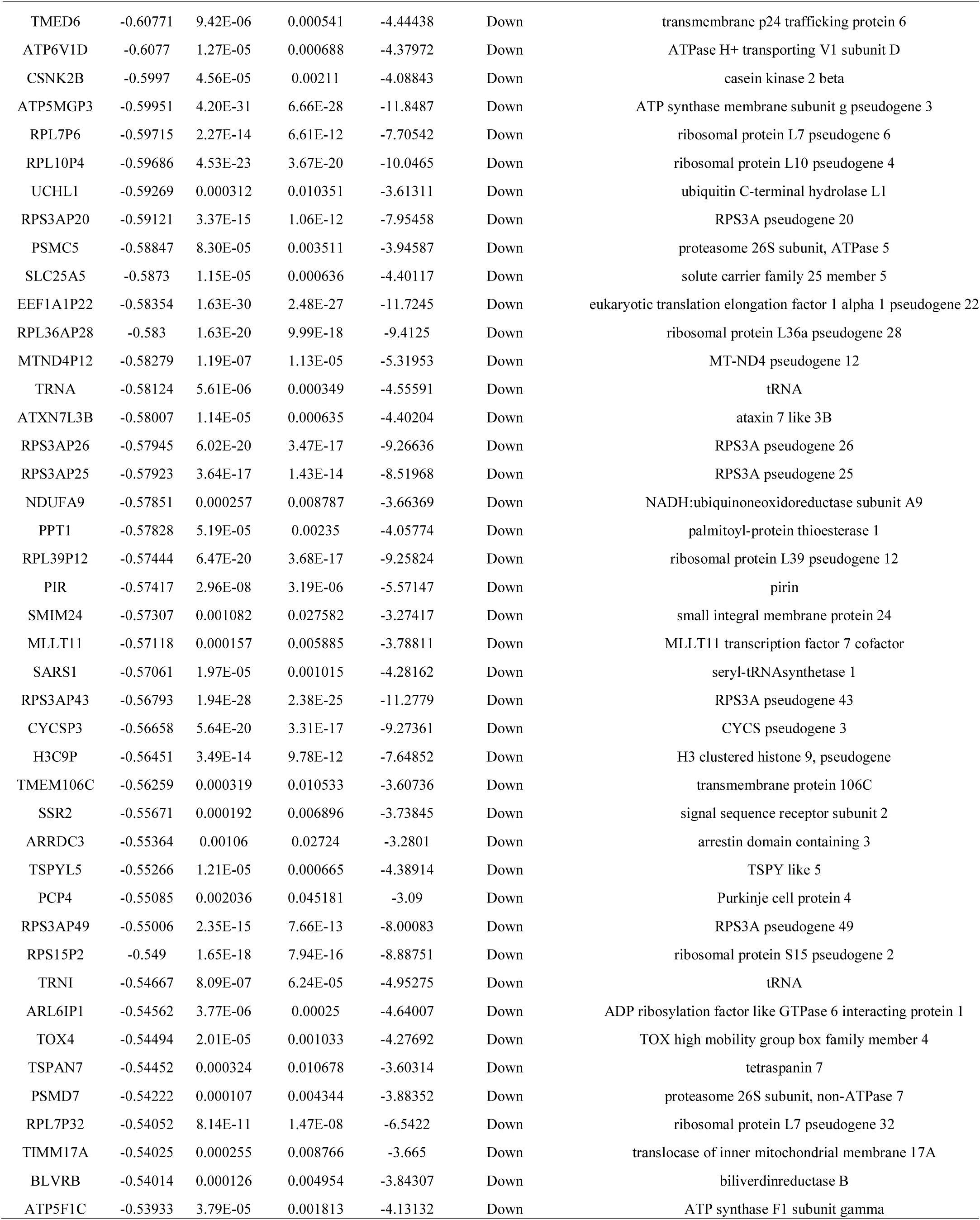
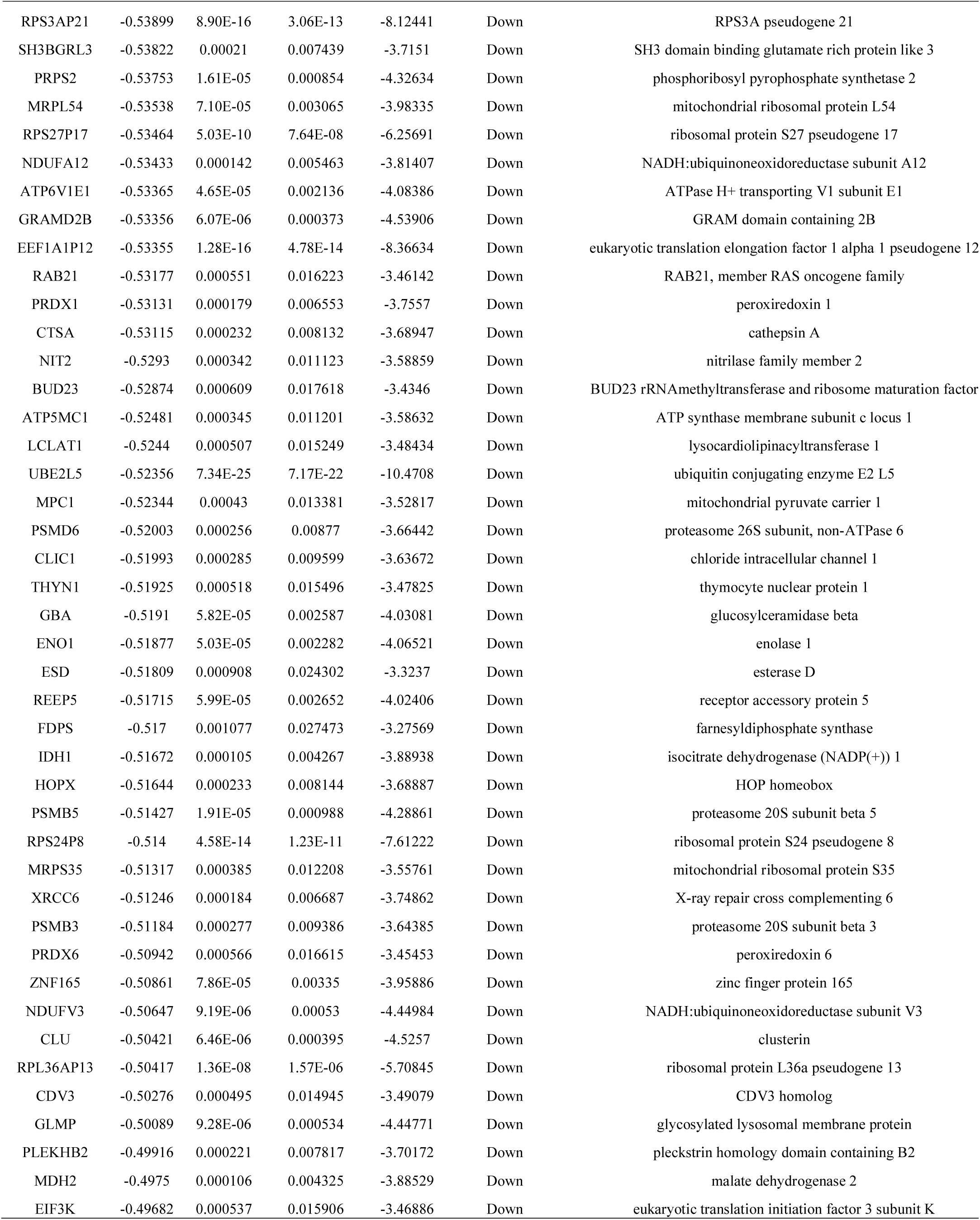
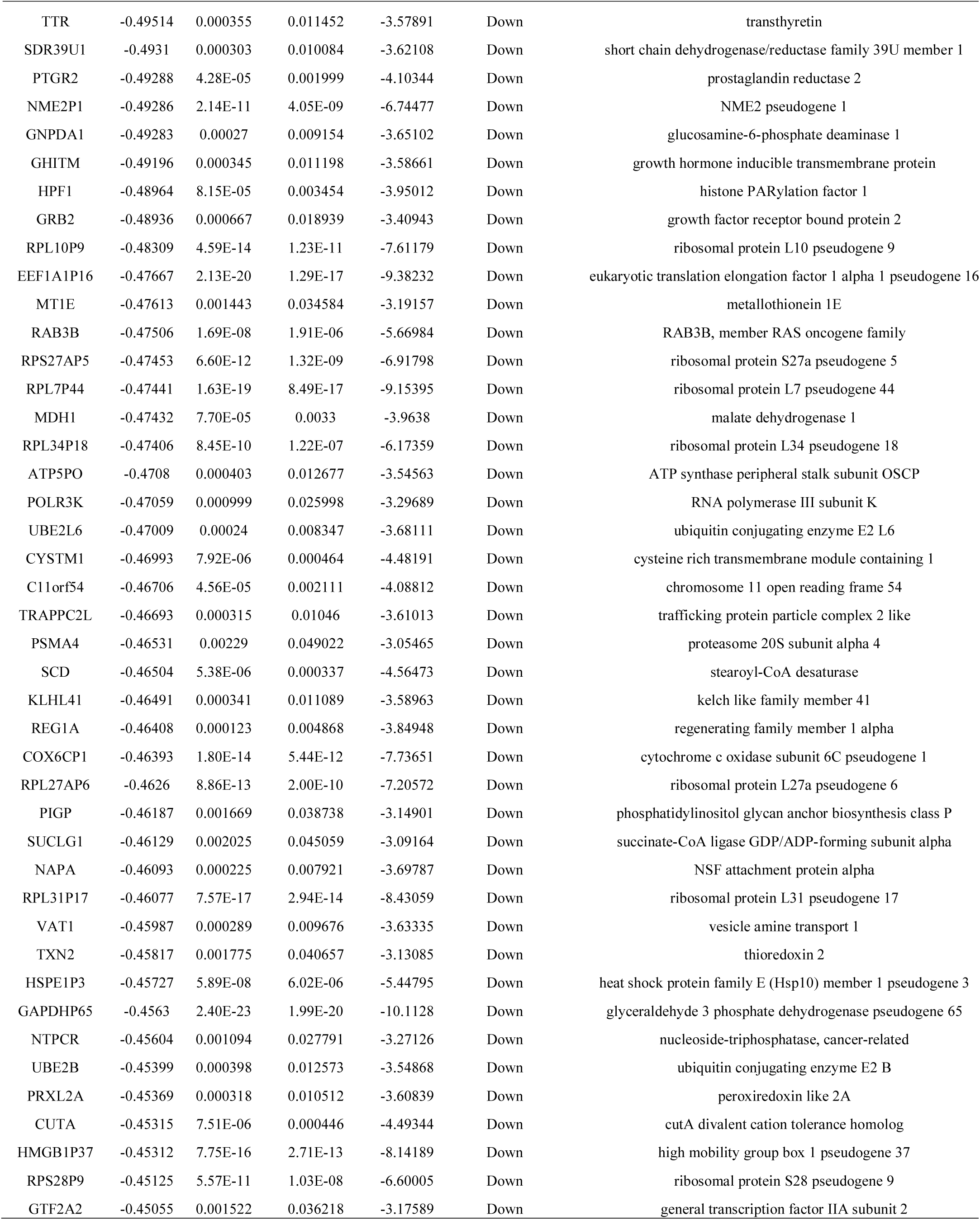
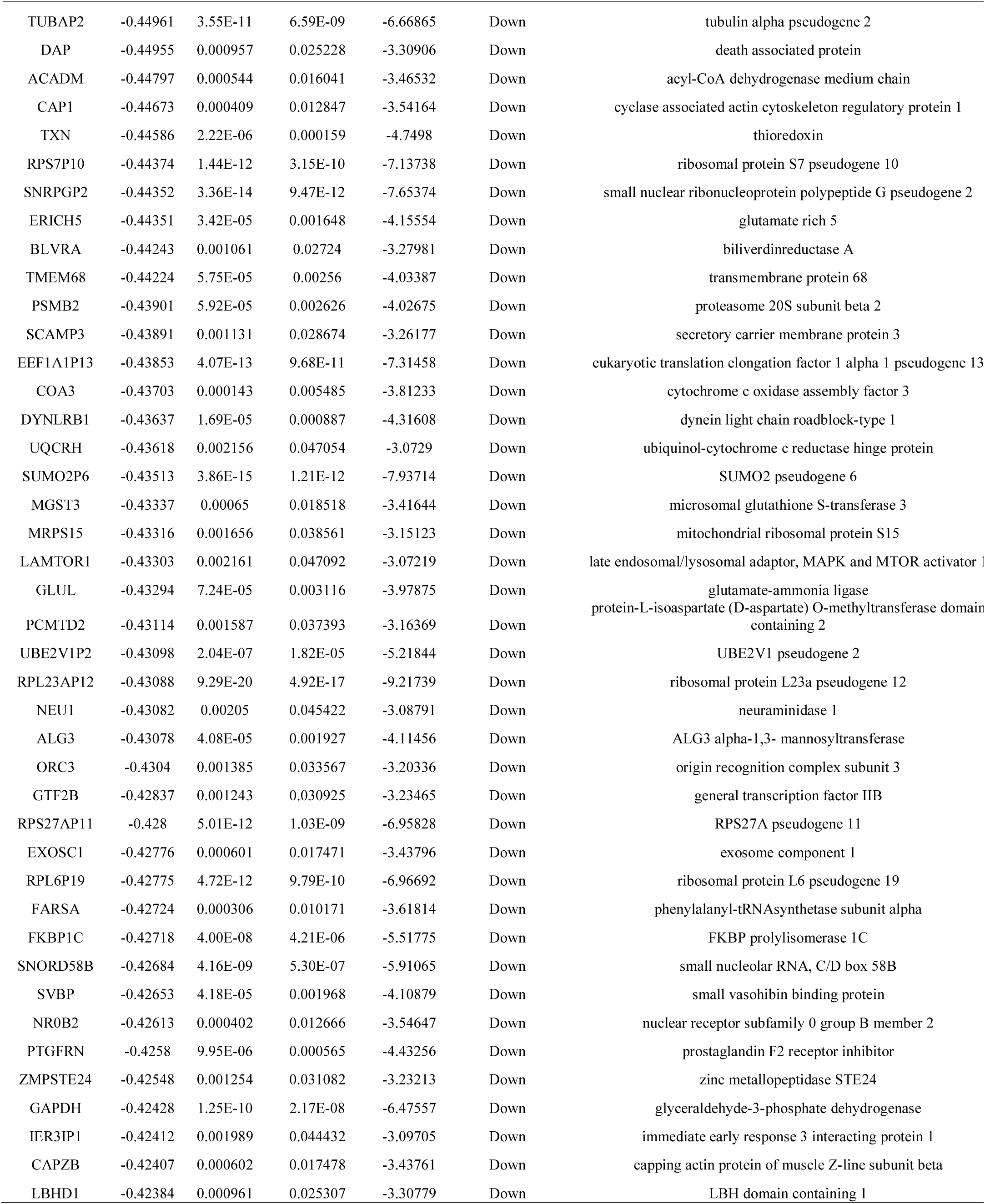
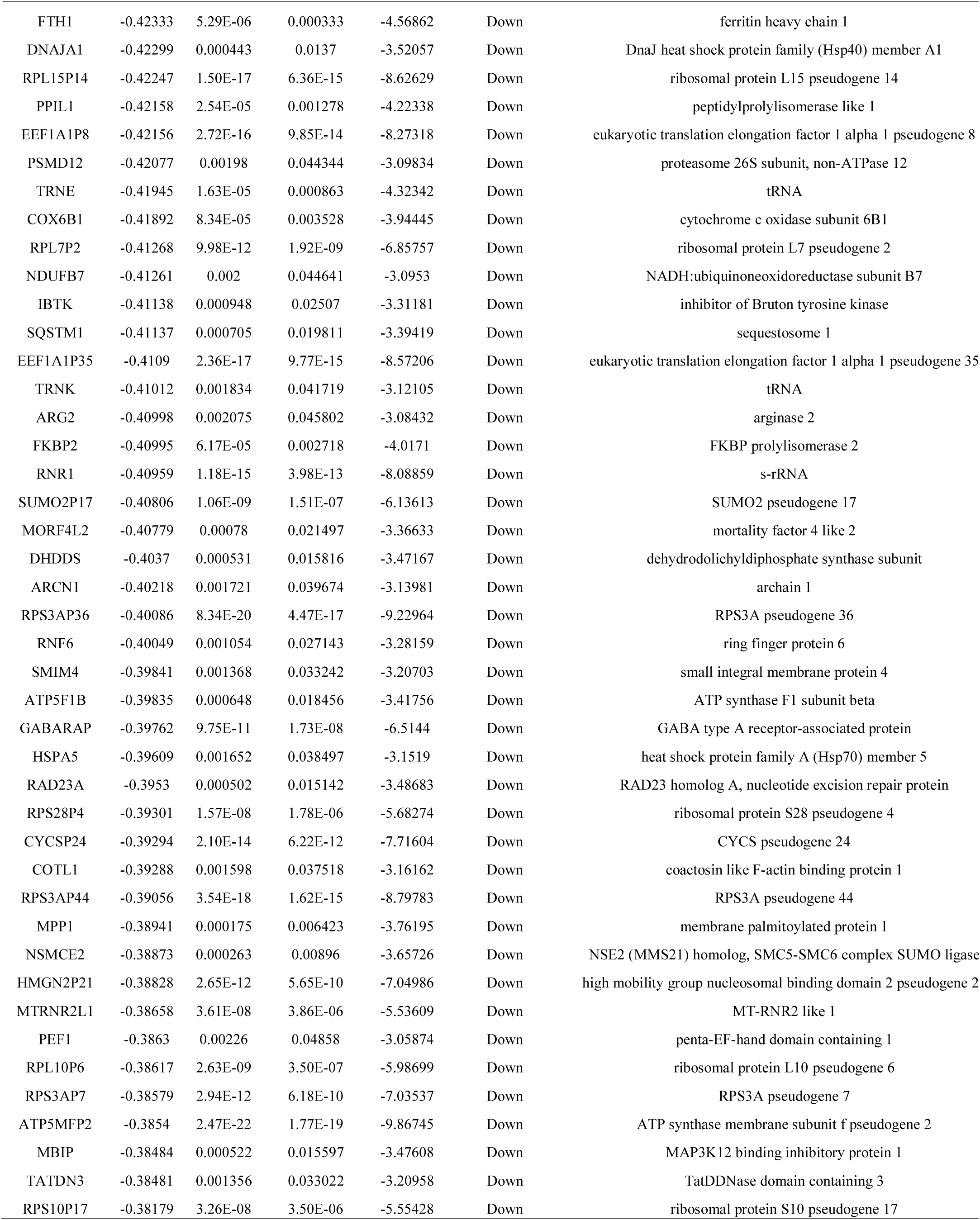
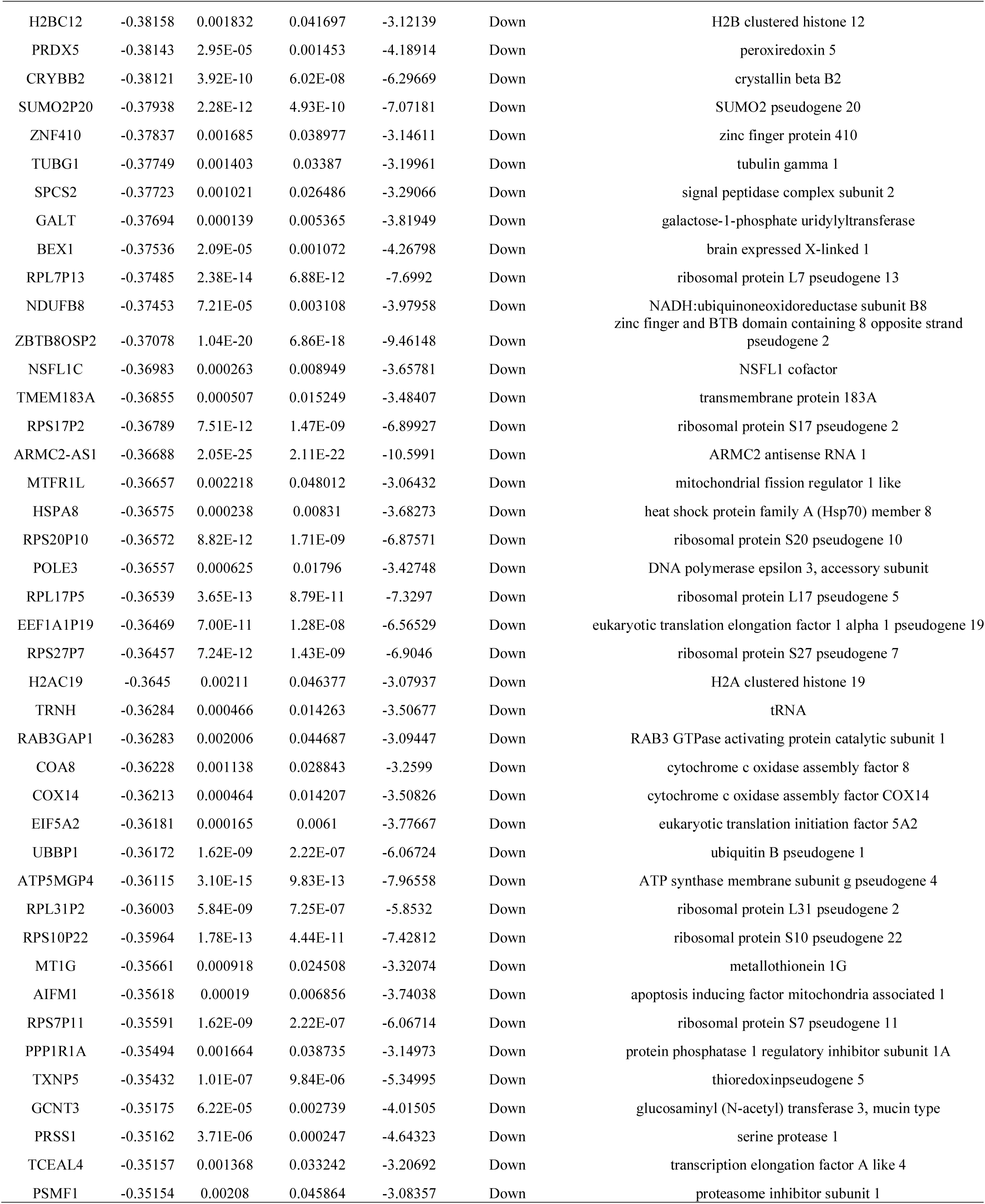
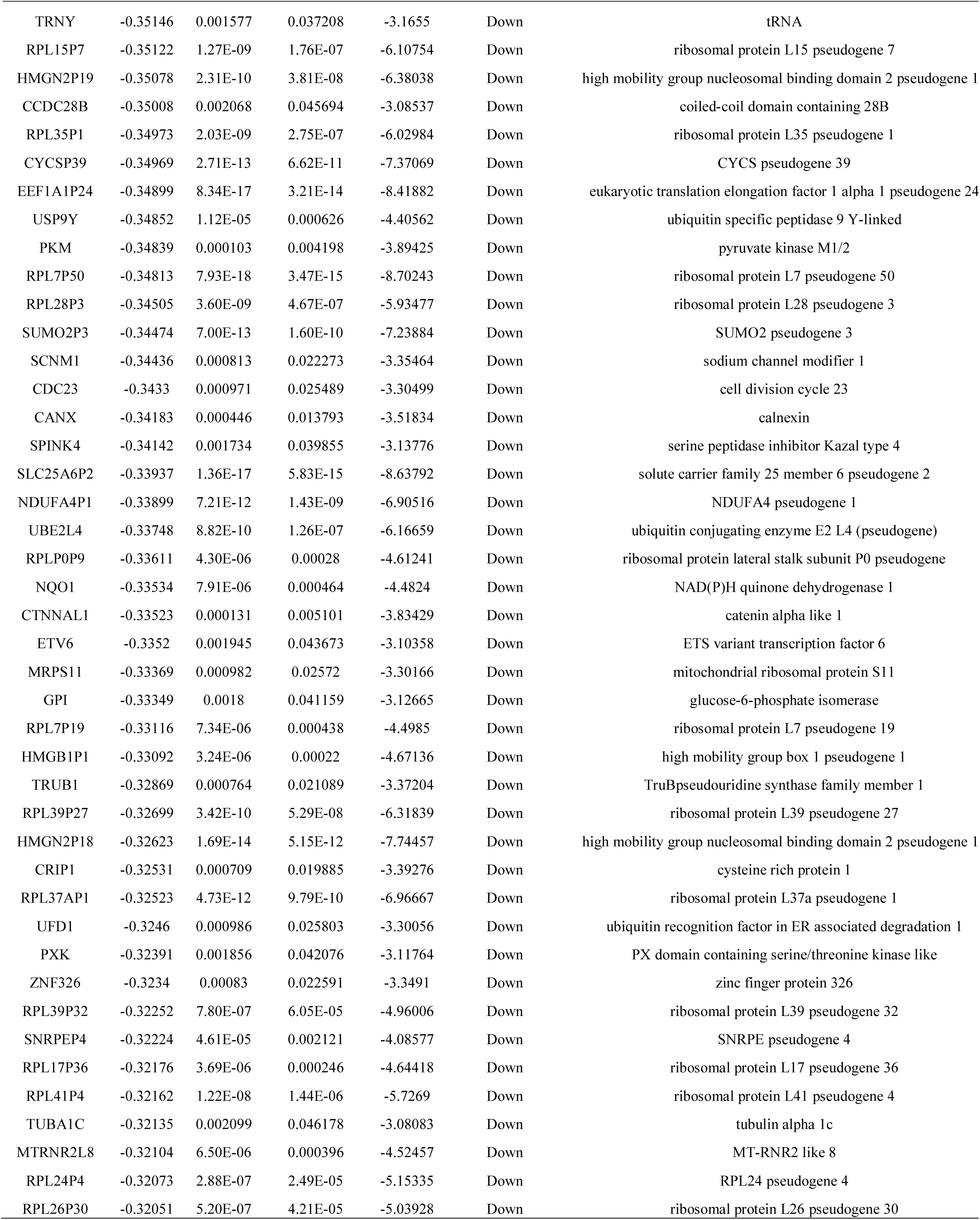
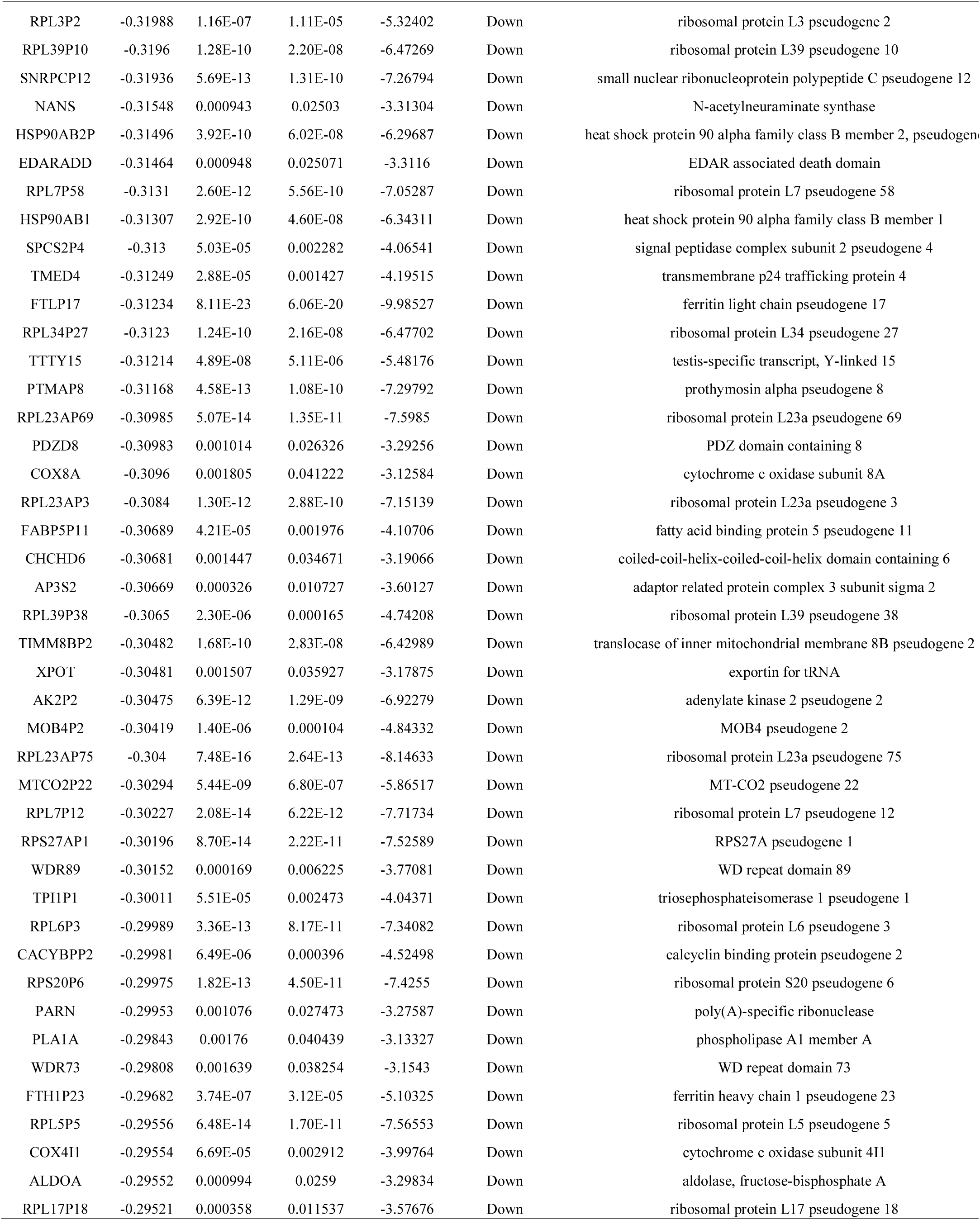
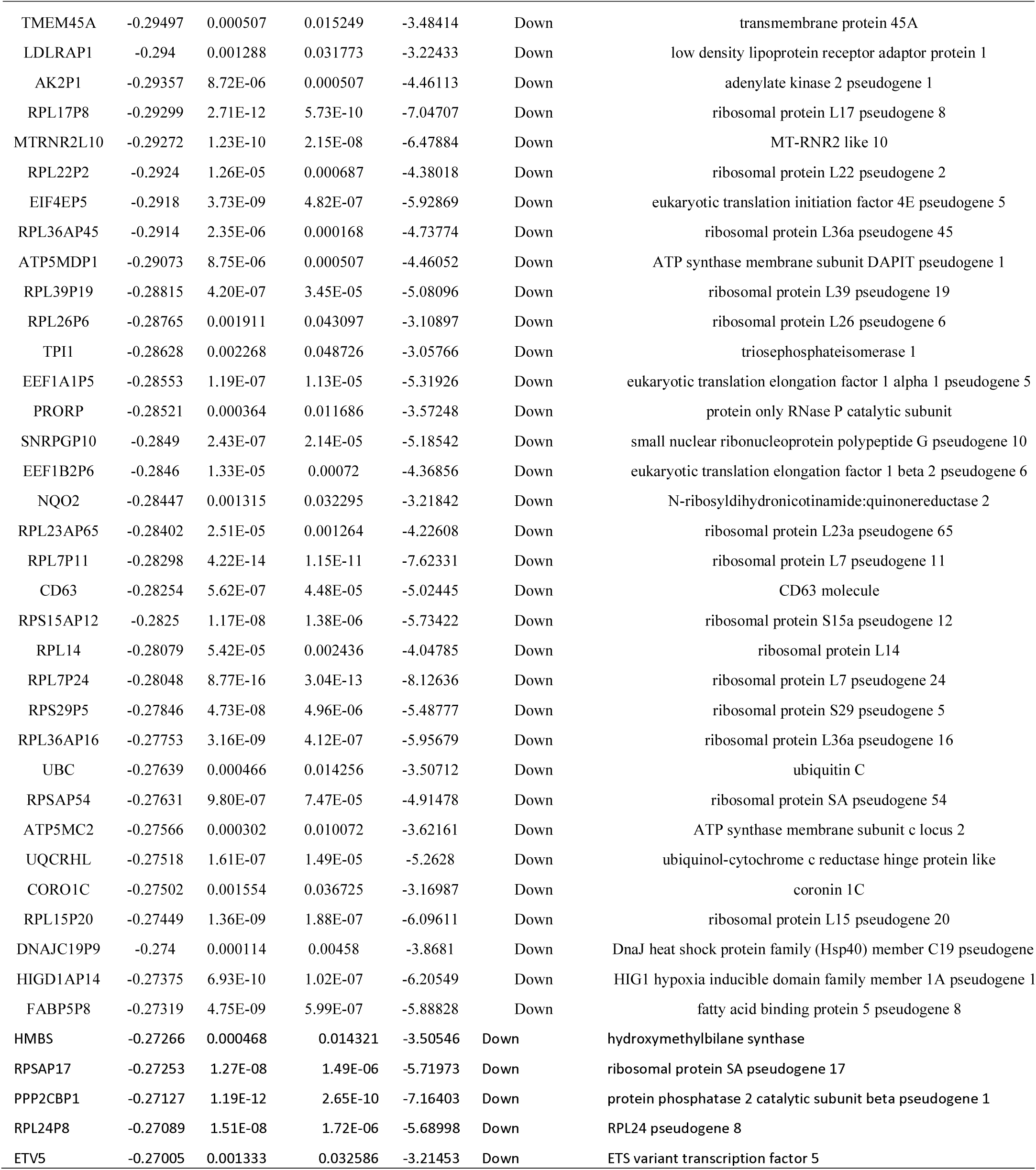
The statistical metrics for key differentially expressed genes (DEGs)

**Fig. 1.**
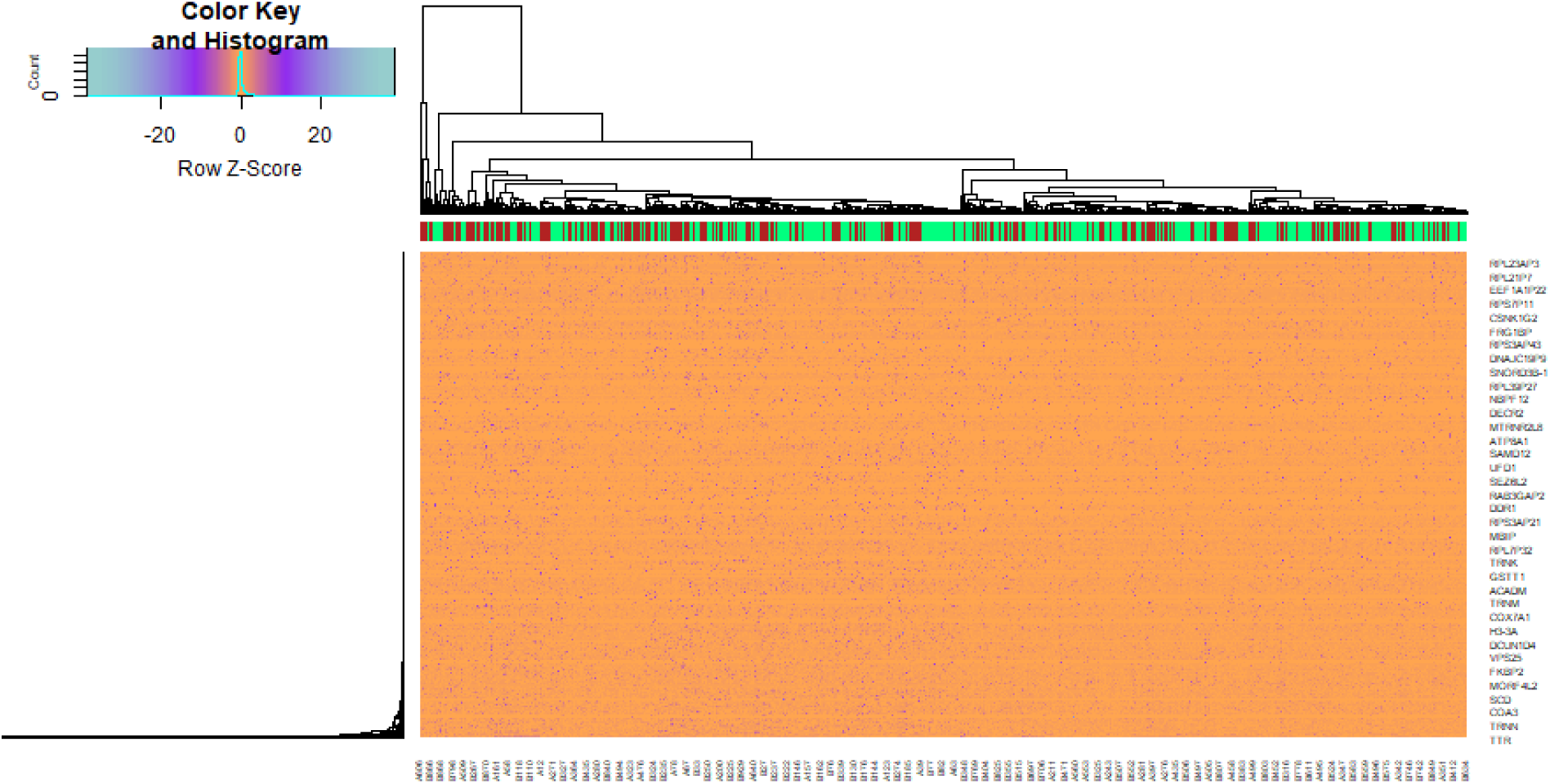
Heat map of differentially expressed genes. Legend on the top left indicate log fold change of genes. (A1 – A651= normal control samples; B1 – B949 = T2DM samples)

**Fig. 2.**
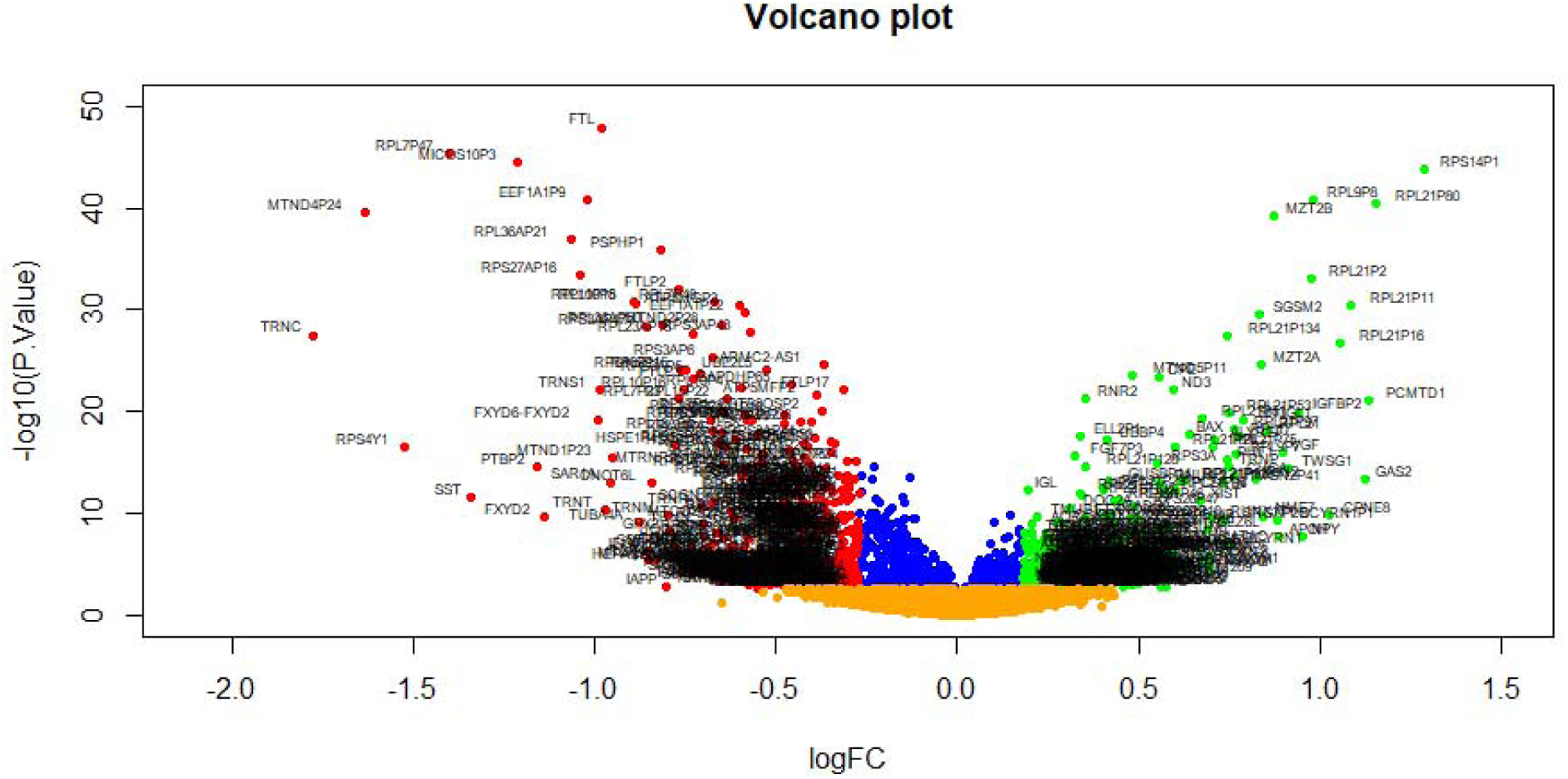
Volcano plot of differentially expressed genes. Genes with a significant change of more than two-fold were selected. Green dot represented up regulated significant genes and red dot represented down regulated significant genes.

### GO and REACTOME pathway enrichment analysis of DEGs

To identify the pathways which had the most significant involvement with the genes identified, up regulated and down regulated genes were submitted into g:Profiler for GO and REACTOME pathway enrichment analysis and are listed in Table 2 and Table 3. GO enrichment analysis revealed that in BP terms, the up regulated genes were mainly enriched in protein metabolic process and positive regulation of biological process. Down regulated genes were mainly enriched in establishment of localization and cellular metabolic process. In CC terms, up regulated genes were mainly enriched in intracellular anatomical structure and endomembrane system, whereas down regulated genes were mainly enriched in cytoplasm and intracellular anatomical structure. In MF terms, up regulated genes were mainly enriched in heterocyclic compound binding and protein binding, whereas down regulated genes were mainly enriched in catalytic activity and protein binding. REACTOME pathway enrichment analysis demonstrated that up regulated genes were significantly enriched in metabolism of proteins and NR1H3 & NR1H2 regulate gene expression linked to cholesterol transport and efflux. Down regulated genes were significantly enriched in the metabolism and the citric acid (TCA) cycle and respiratory electron transport.

**Table 2.**
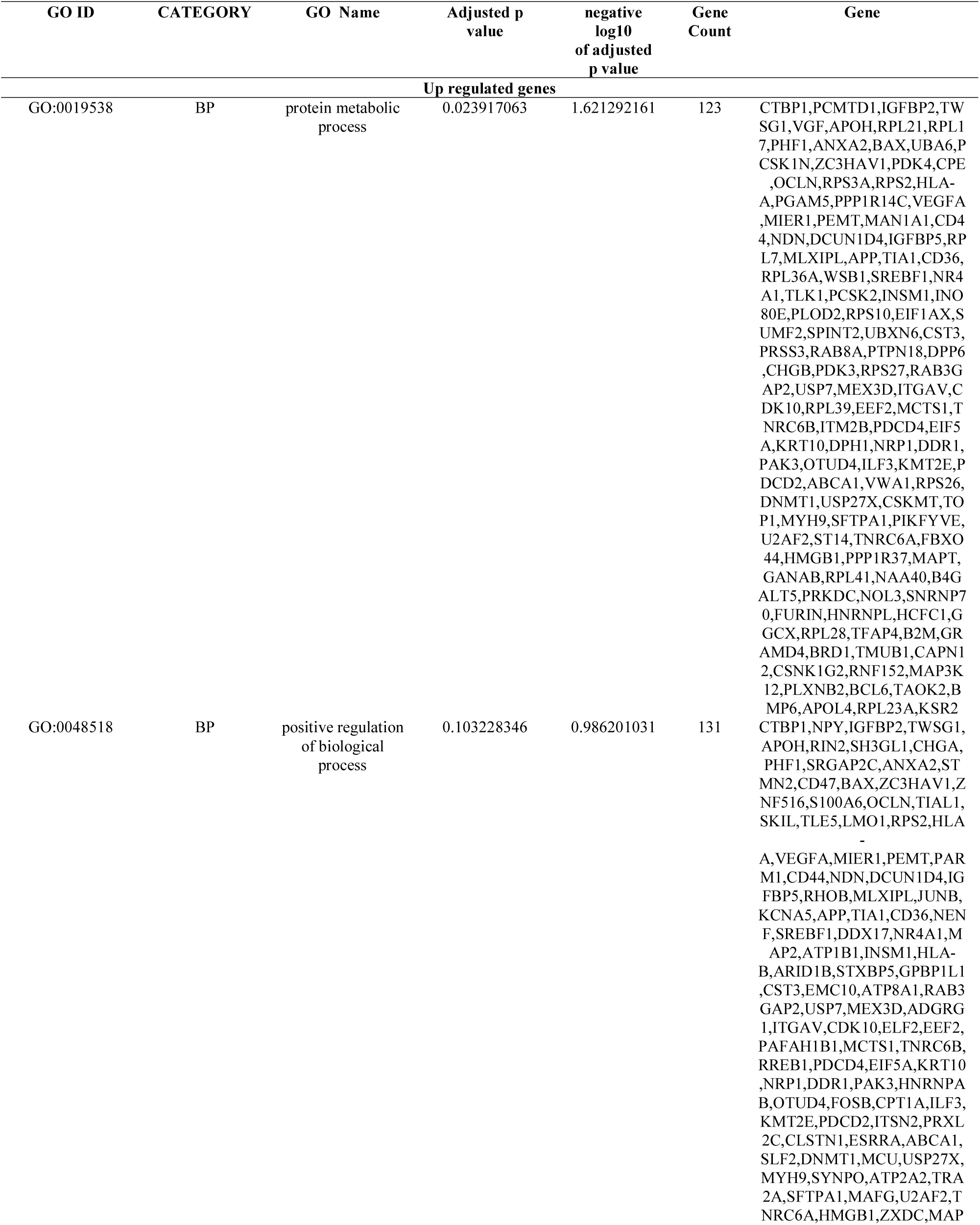
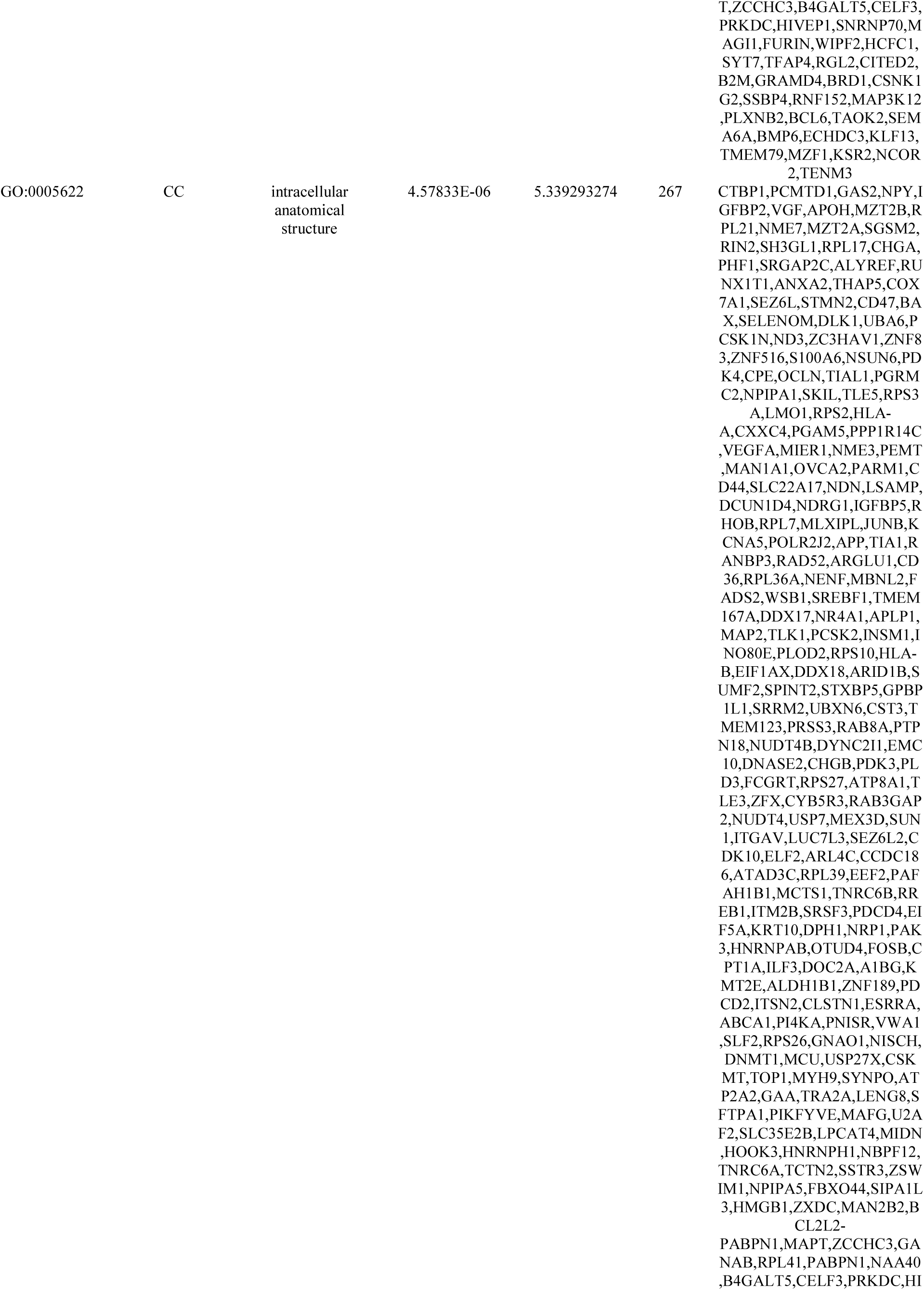
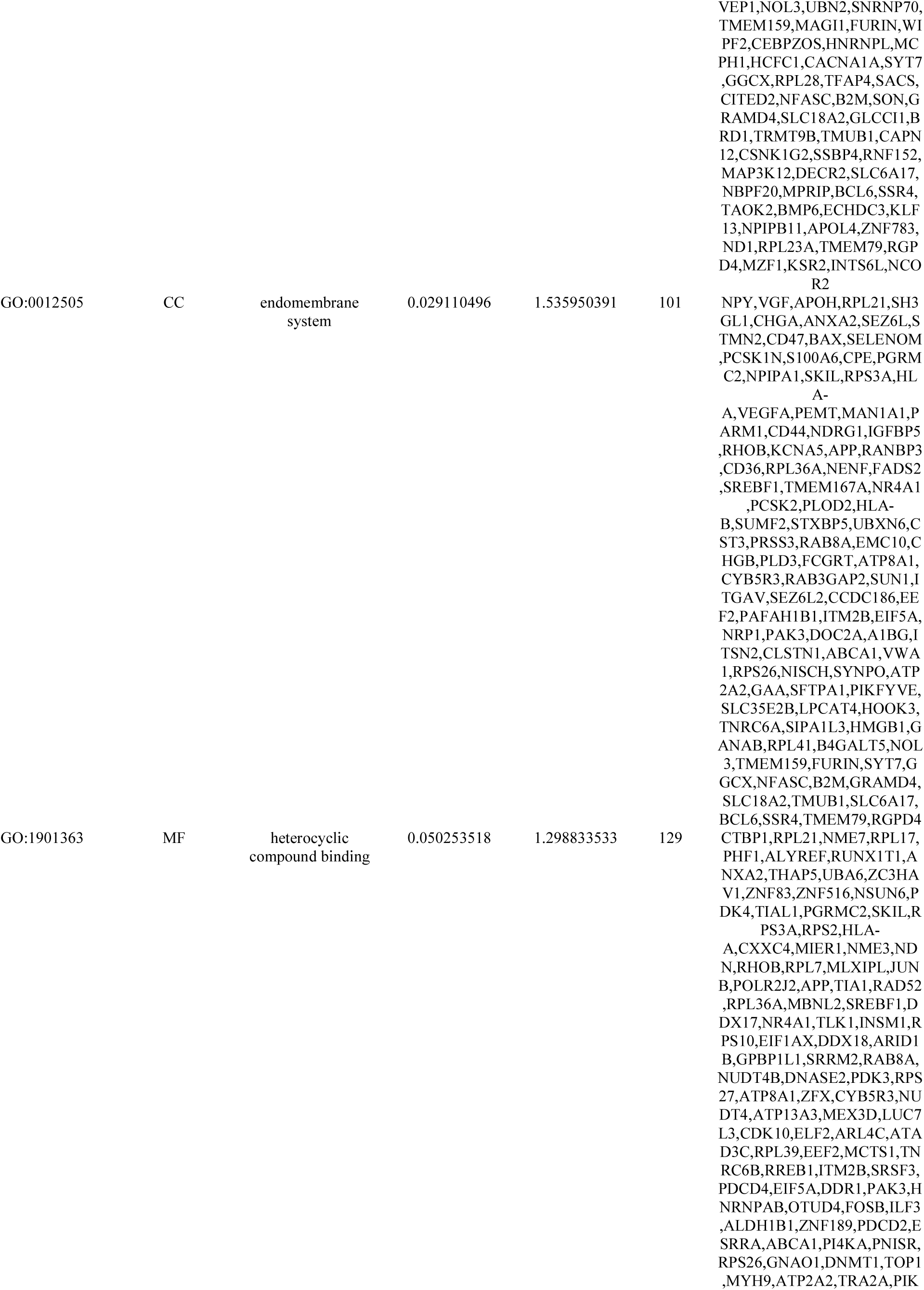
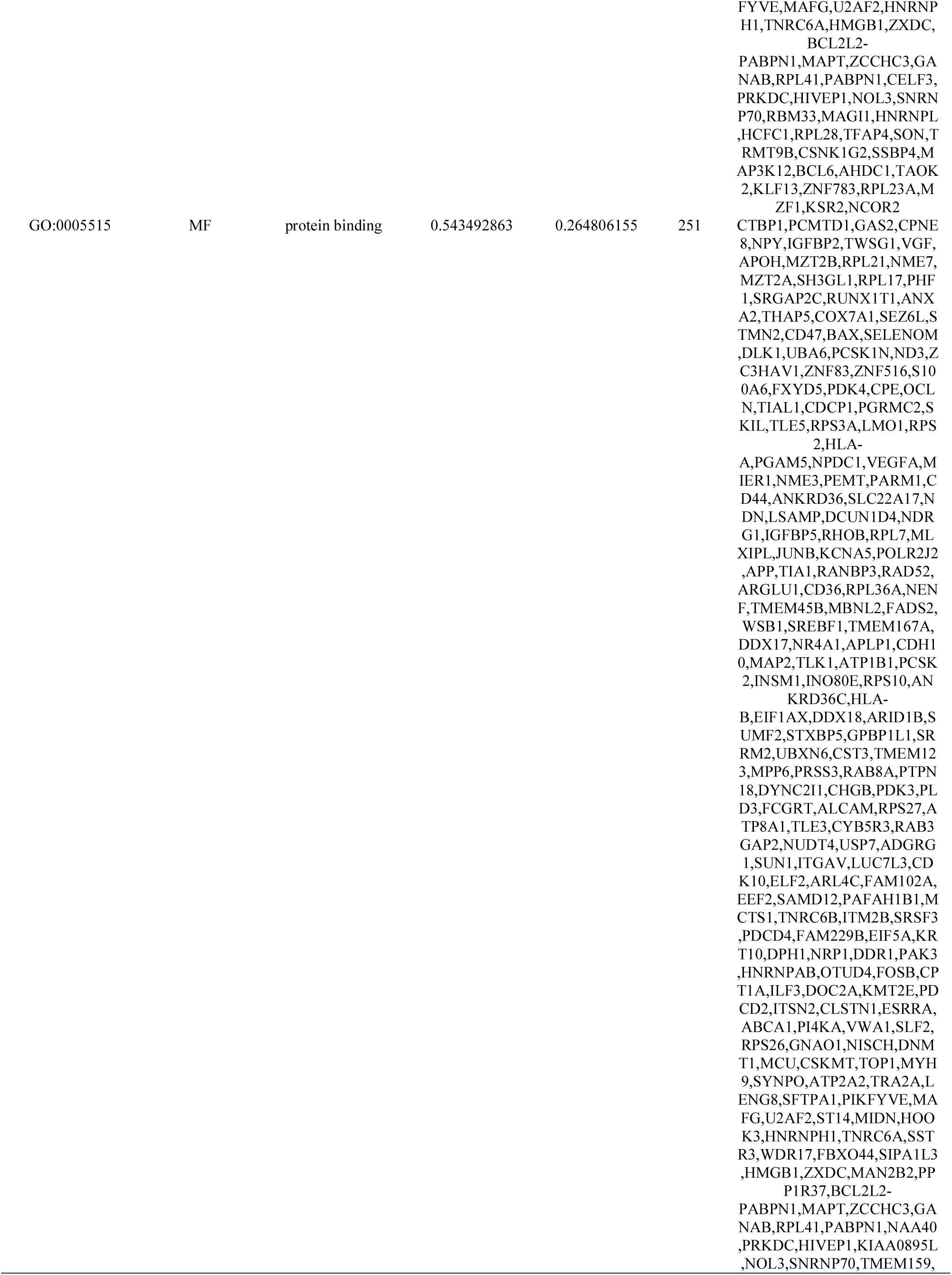
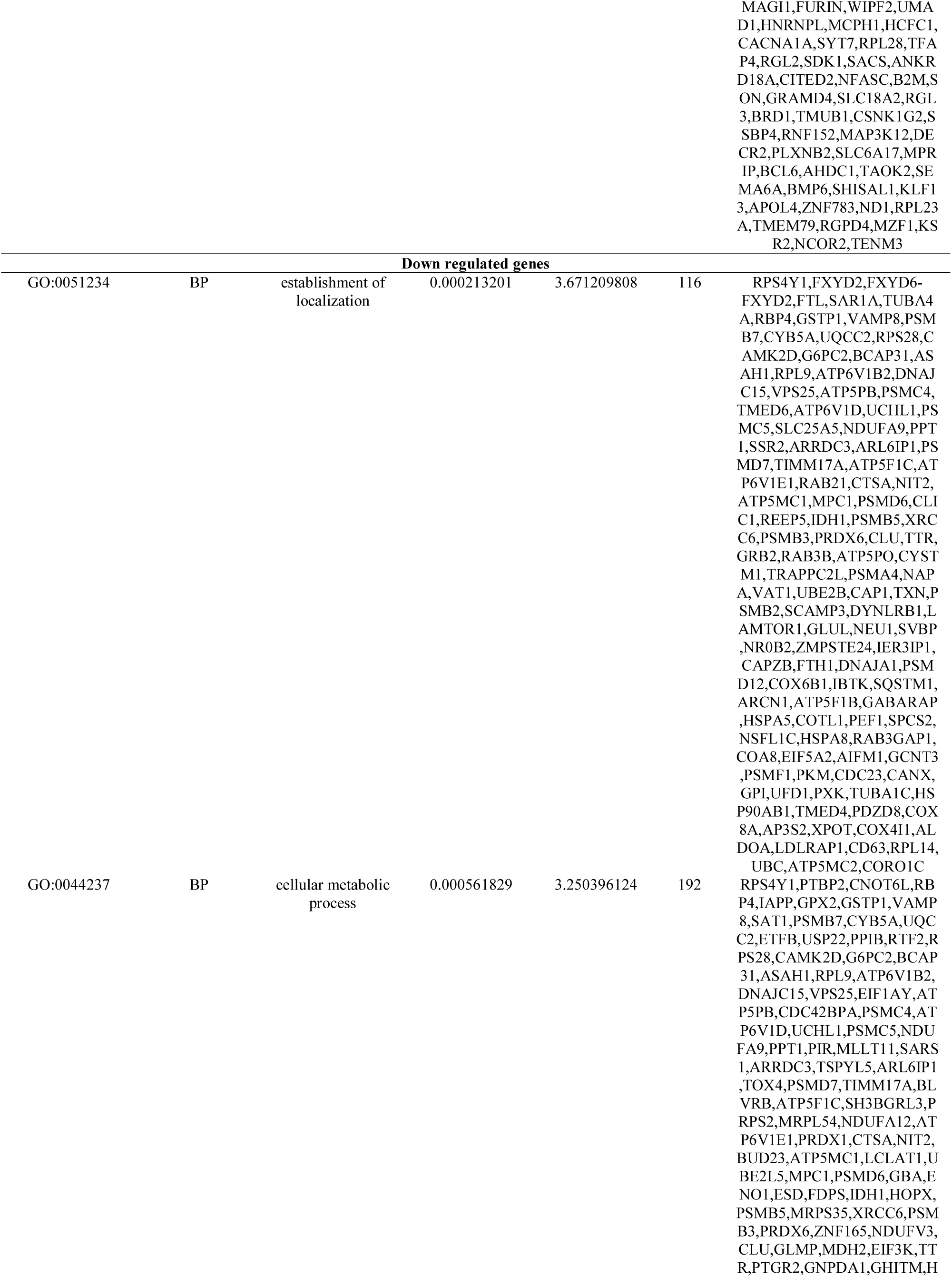
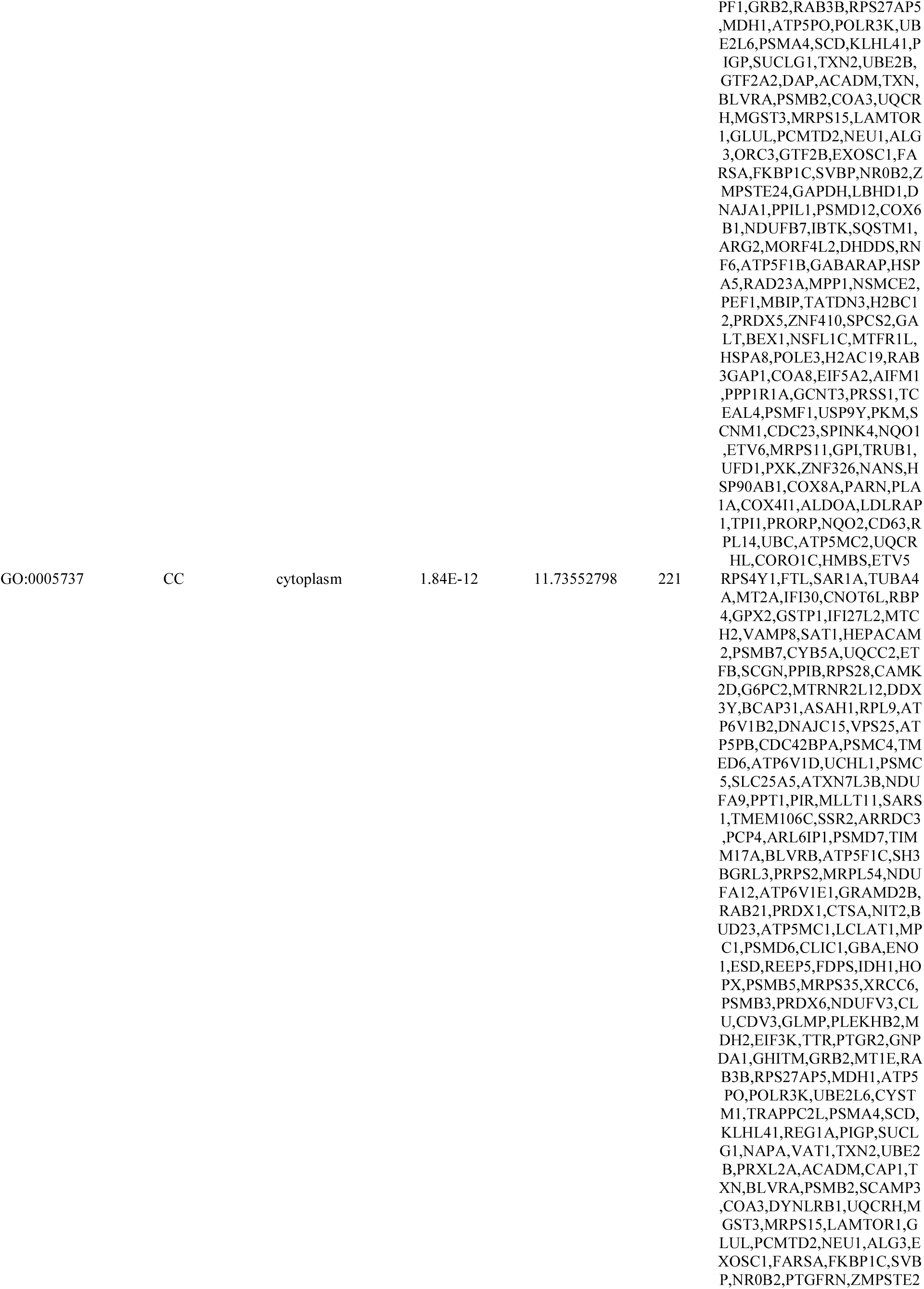
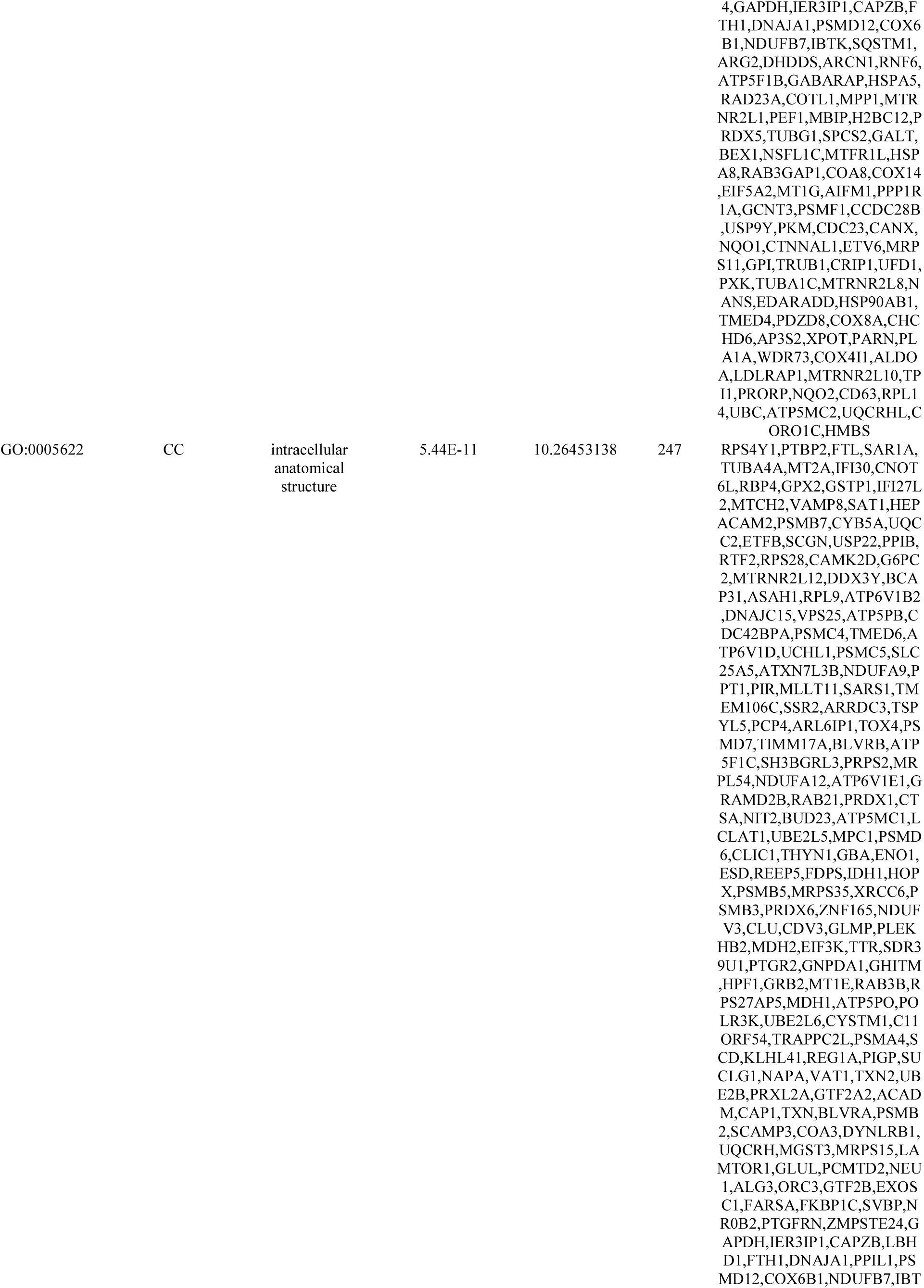
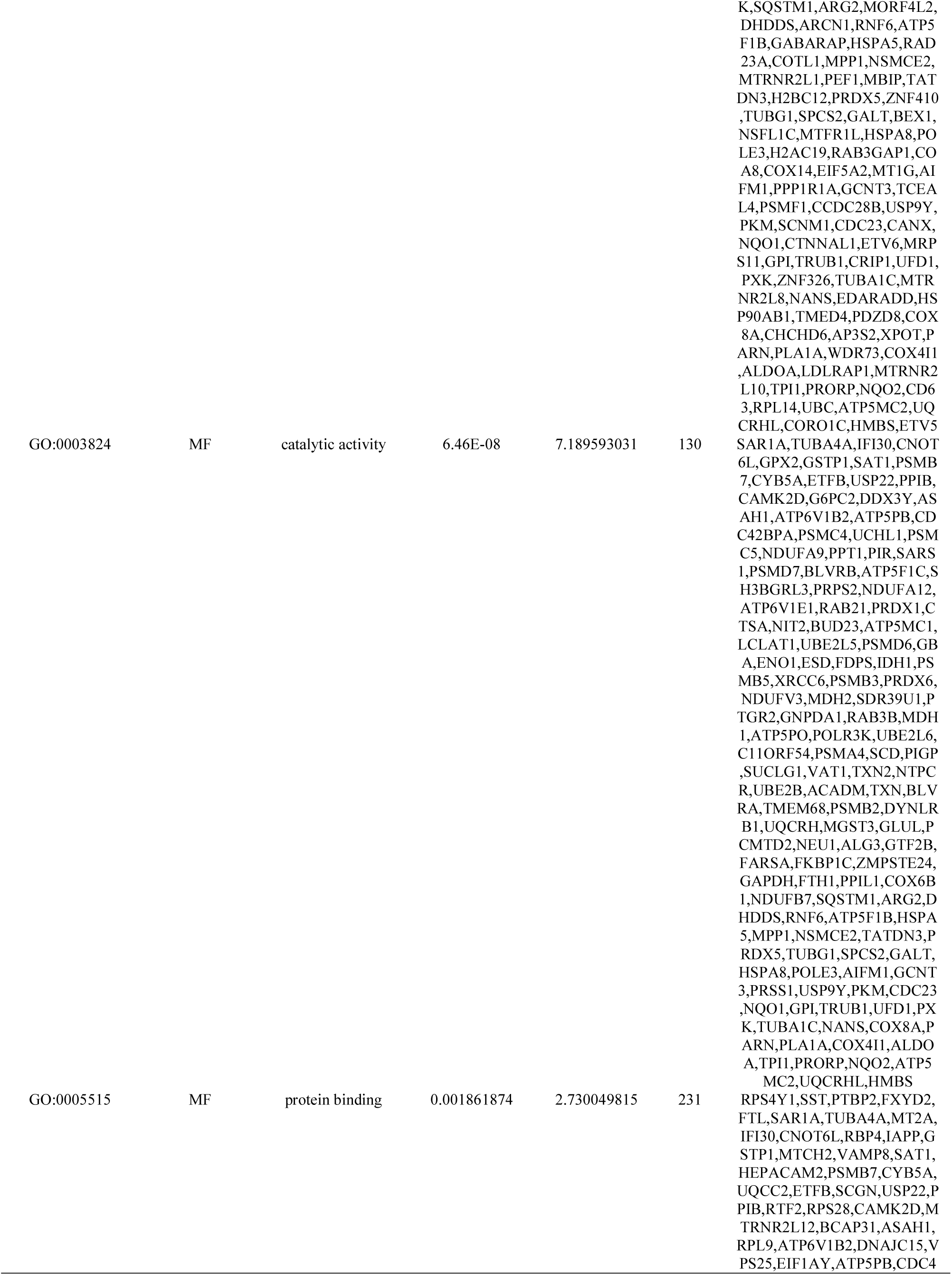
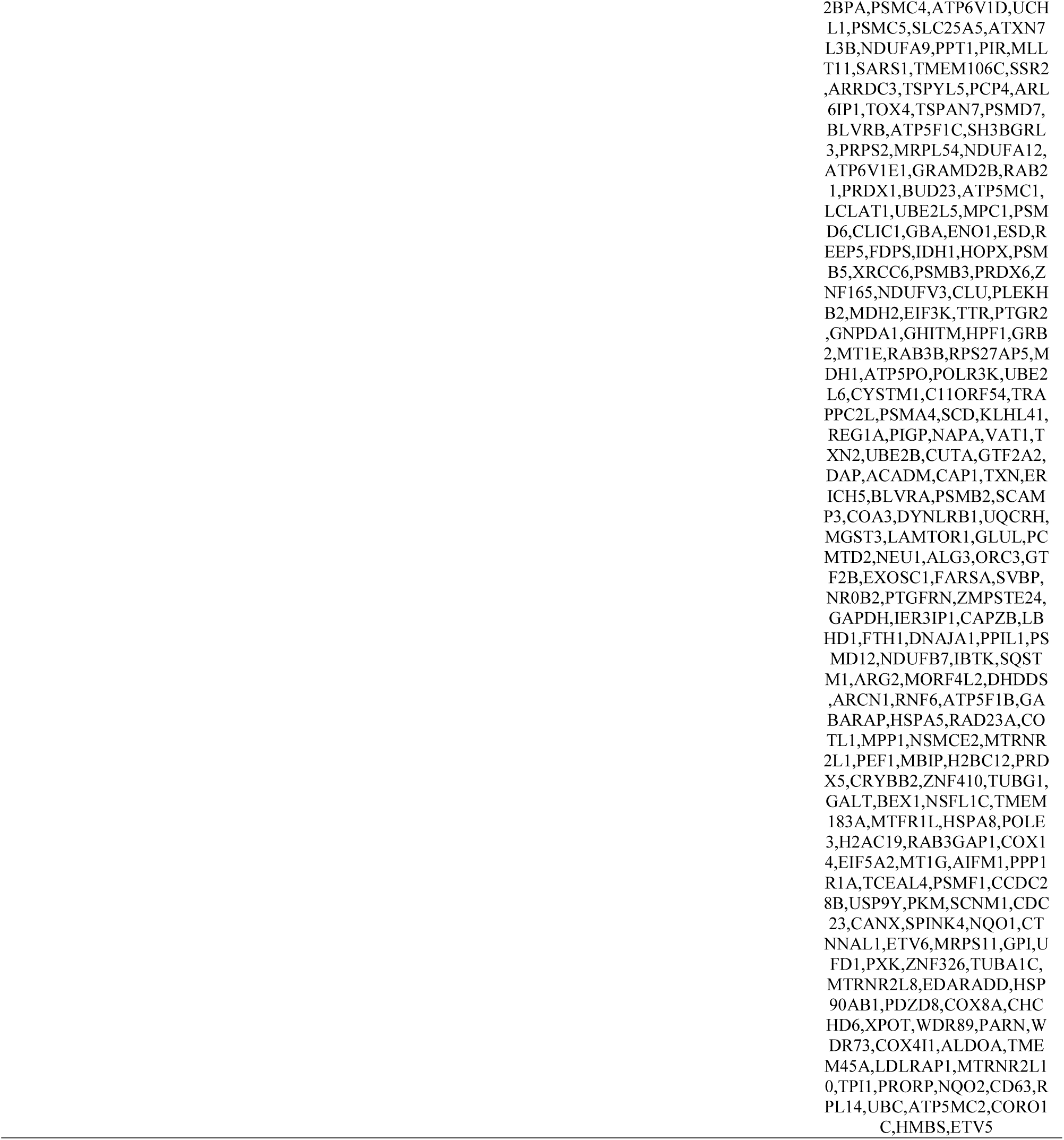
The enriched GO terms of the up and down regulated differentially expressed genes

**Table 3.**
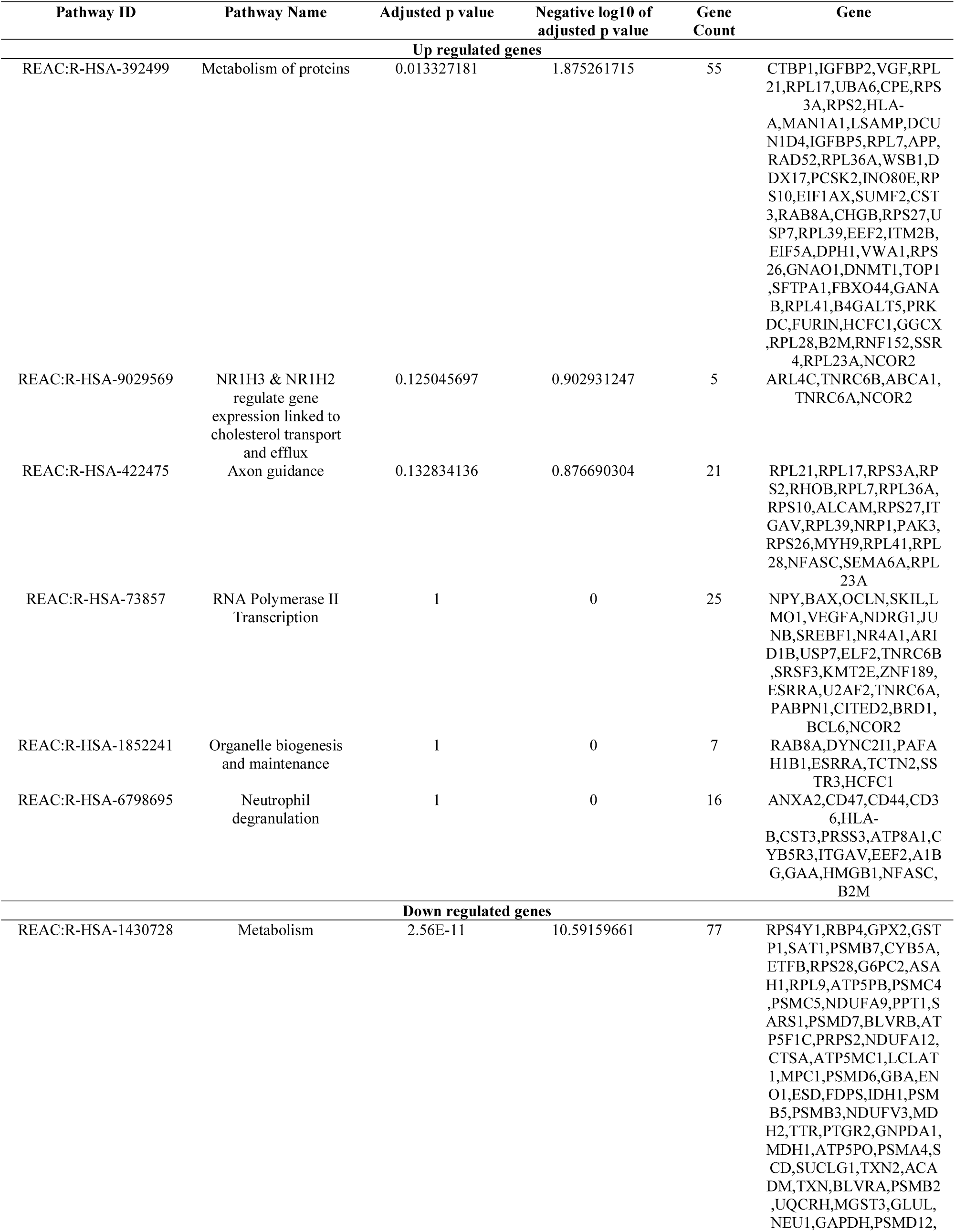
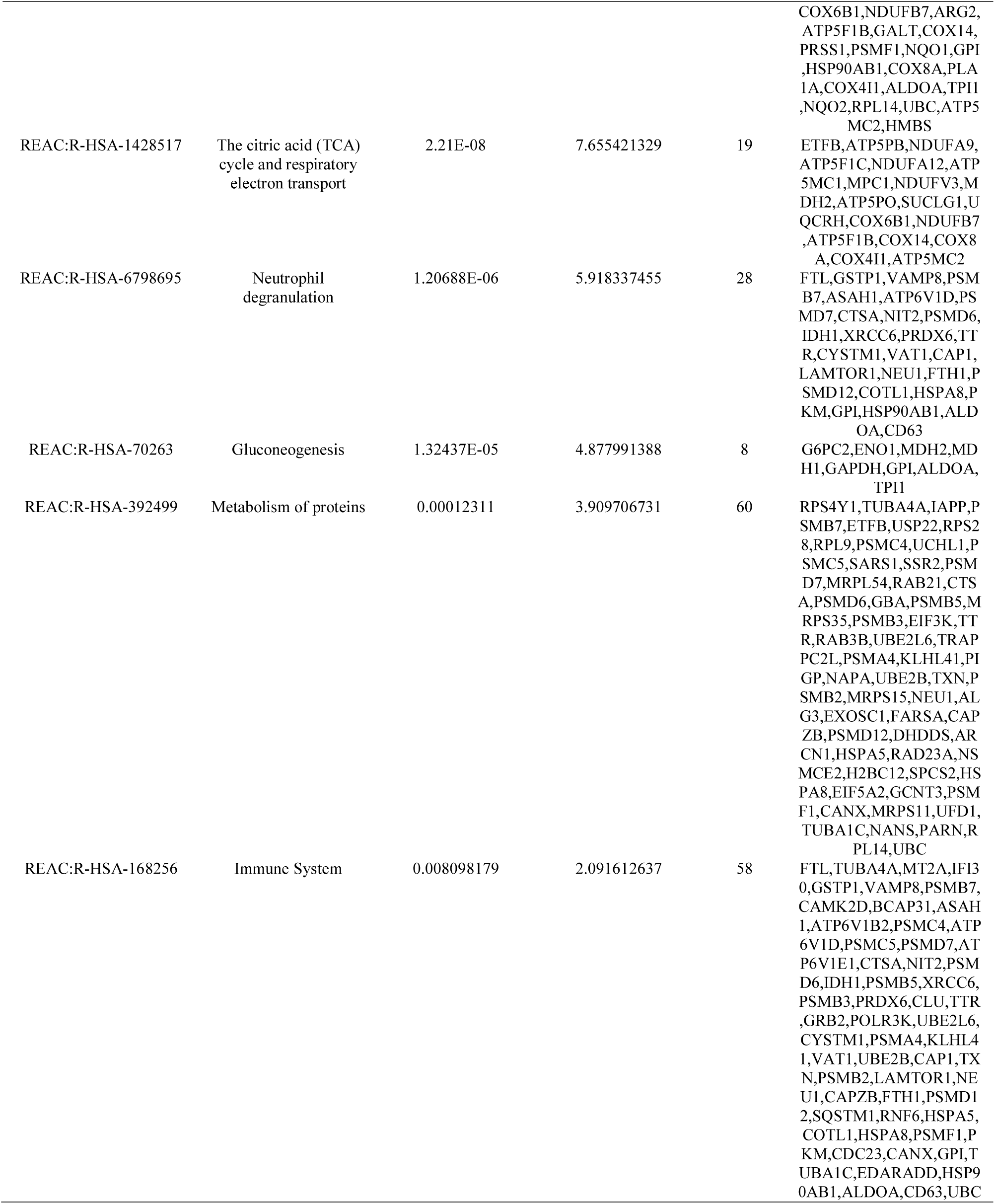
The enriched pathway terms of the up and down regulated differentially expressed genes

### Construction of the PPI network and module analysis

Following the analysis based on the PPI networks, 4424 nodes and 8670 edges were identified in Cytoscape (Fig. 3). The genes with higher scores were the hub genes, as the genes of higher node degree, betweenness centrality, stress centrality and closeness centrality might be linked with T2DM. The top hub genes were APP, MYH9, TCTN2, USP7, SYNPO, GRB2, HSP90AB1, UBC, HSPA5 and SQSTM1, and are listed in Table 4. A total of two modules were selected through PEWCC1 analysis, and module 1 had nodes 98 and edges 117 (Fig. 4A) and module 2 had nodes 81 and edges 248 (Fig. 4B). Enrichment analysis demonstrated that modules 1 and 2 might be linked with RNA polymerase II transcription, intracellular anatomical structure, metabolism of proteins, protein metabolic process, positive regulation of biological process, metabolism, immune system, establishment of localization, cytoplasm, neutrophil degranulation, cellular metabolic process, intracellular anatomical structure and protein binding.

**Fig. 3.**
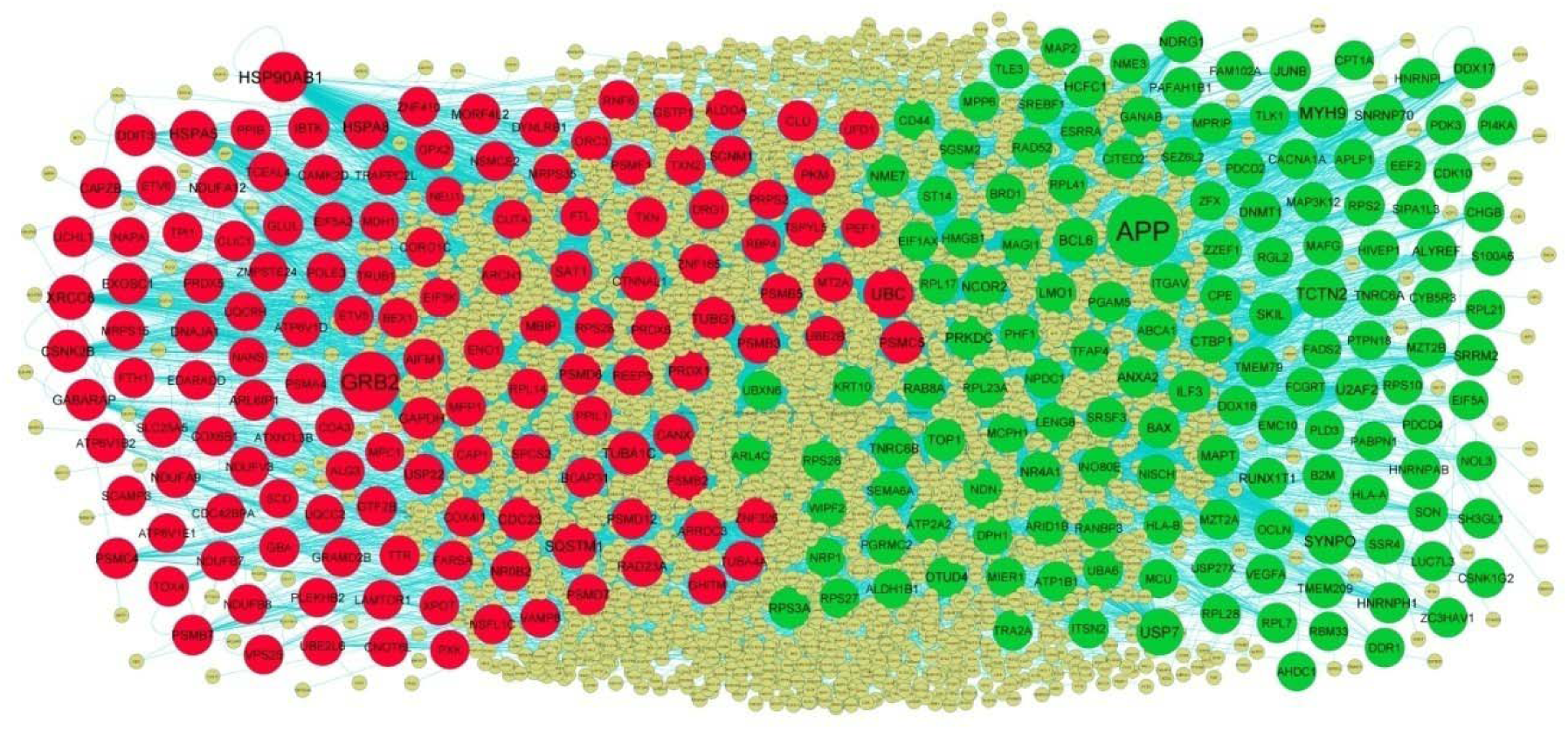
PPI network of DEGs. The PPI network of DEGs was constructed using Cytoscap. Up regulated genes are marked in green; down regulated genes are marked in red

**Fig. 4.**
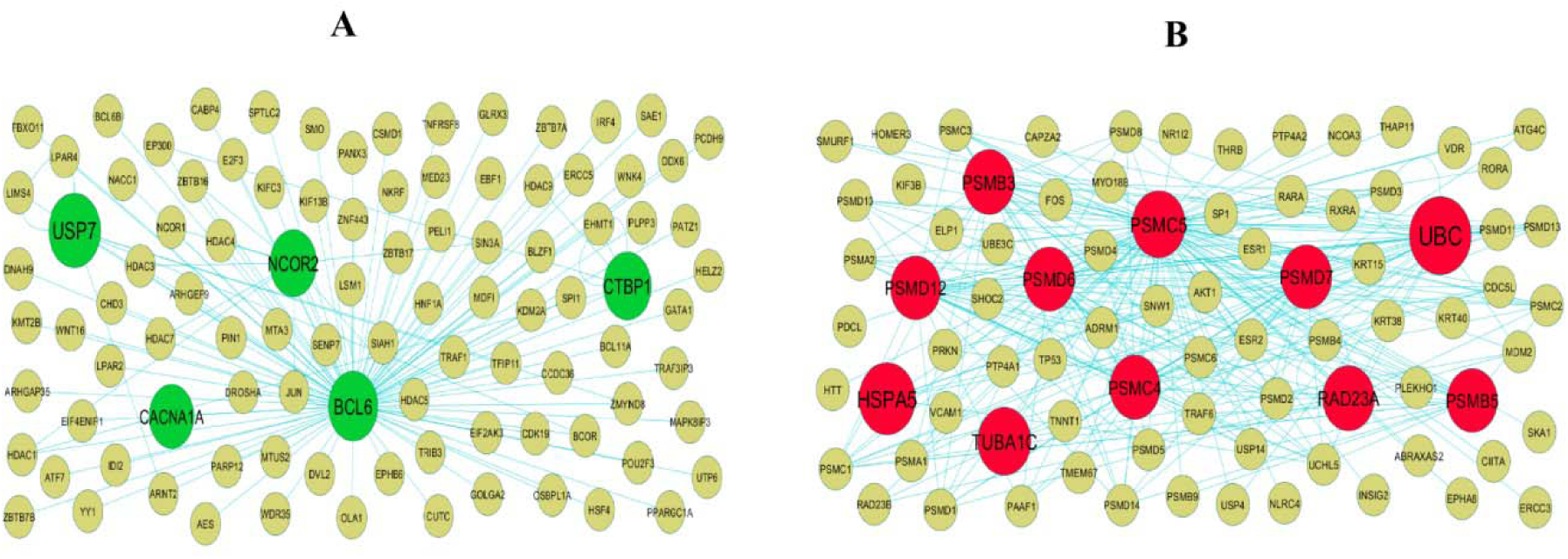
Modules of isolated form PPI of DEGs. (A) The most significant module was obtained from PPI network with 98 nodes and 117 edges for up regulated genes (B) The most significant module was obtained from PPI network with 81 nodes and 248 edges for down regulated genes. Up regulated genes are marked in green; down regulated genes are marked in red.

**Table 4.**
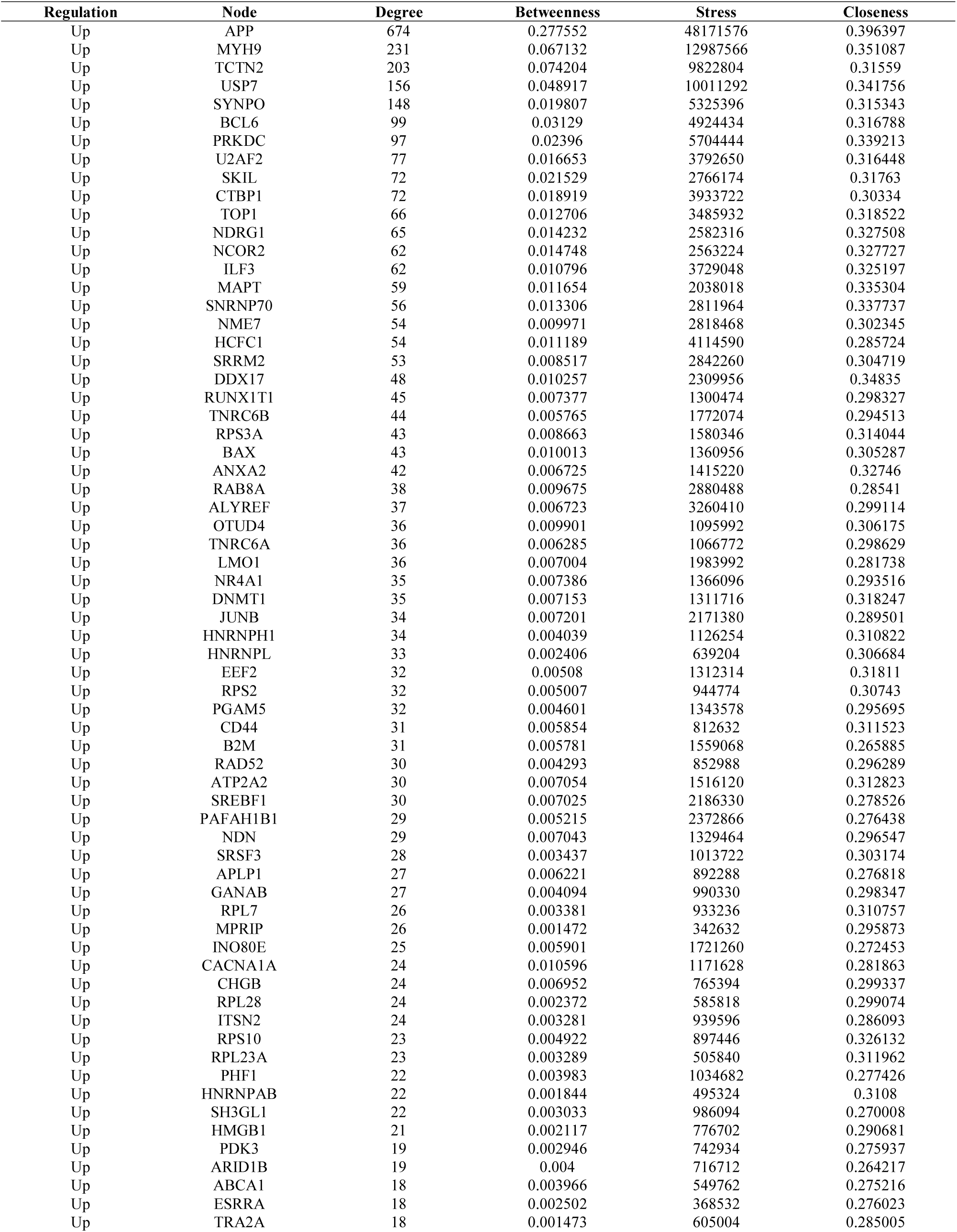
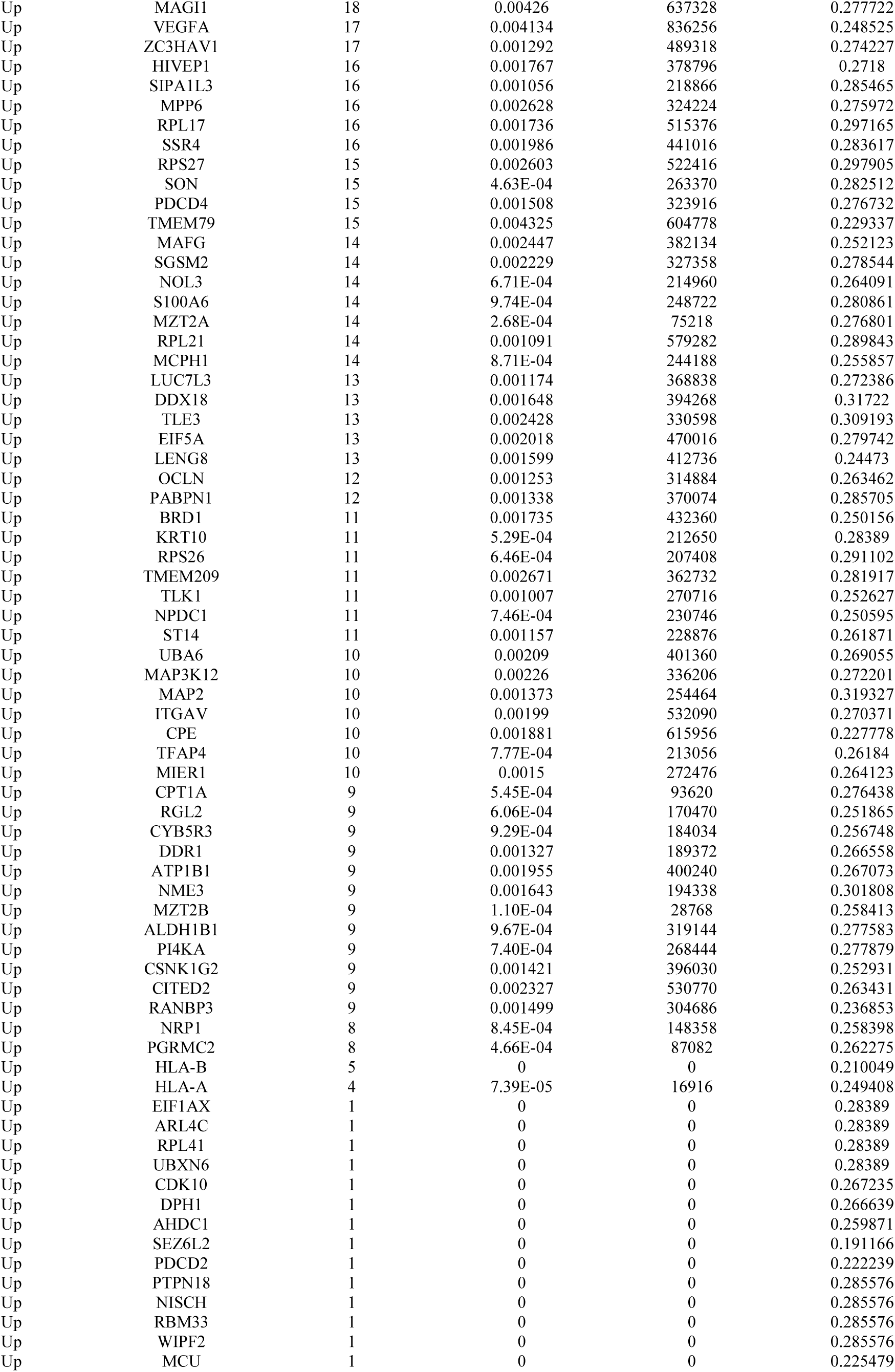
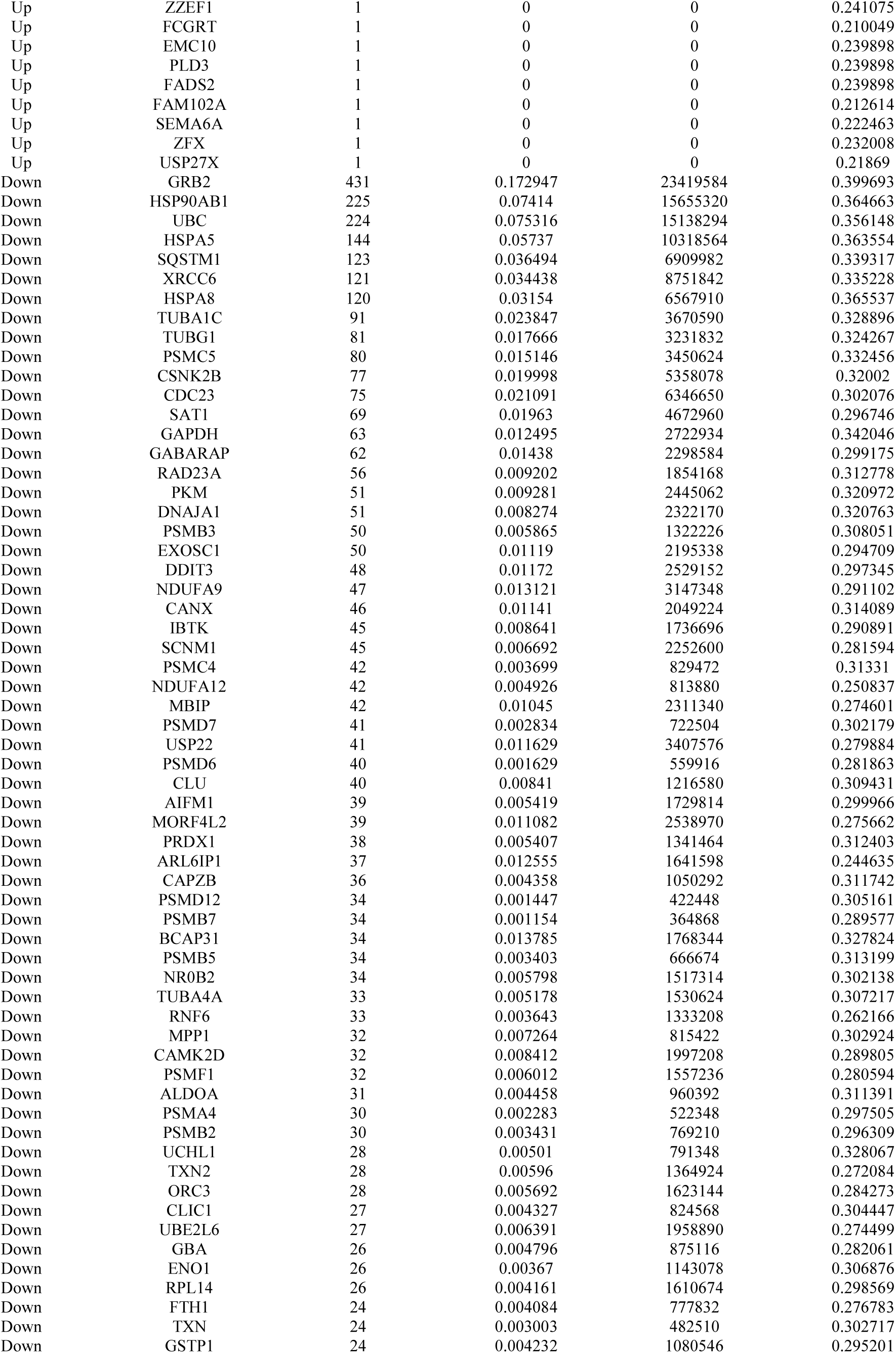
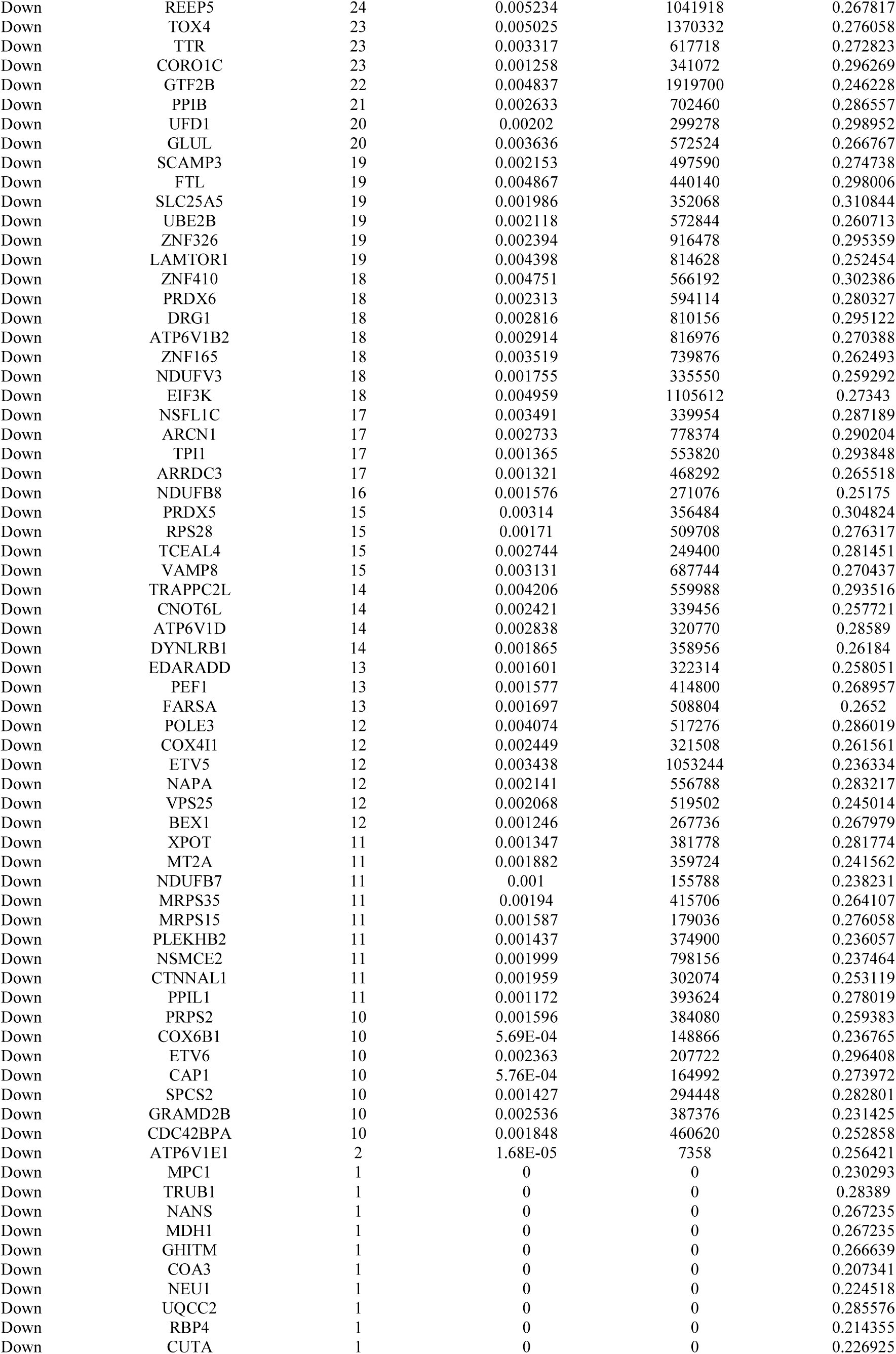
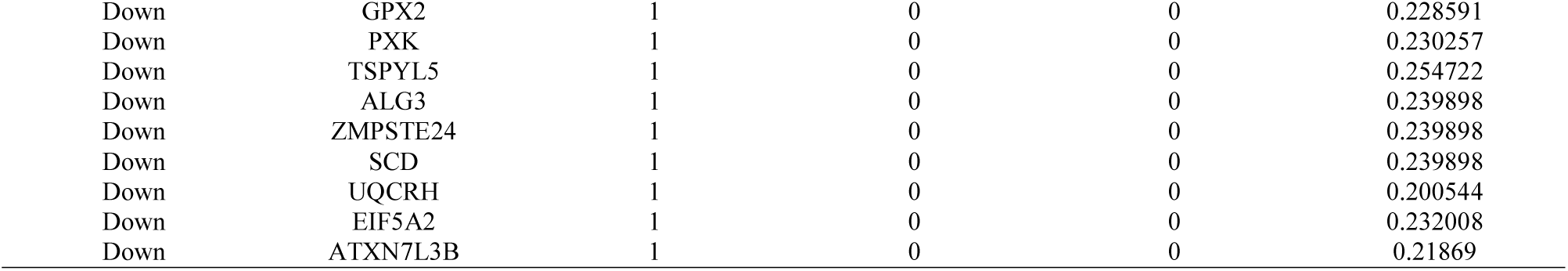
Topology table for up and down regulated genes.

### MiRNA-hub gene regulatory network construction

The hub genes of the DEGs in T2DM were performed by online databases miRNet. Based on the miRNAs, a miRNA -hub gene regulatory network was constructed with 2630 nodes (miRNA: 2345 and hub gene: 285) and 20765 interaction pairs (Fig. 5). PRKDC was the gene targets of 163 miRNAs (ex; hsa-mir-142-5p), MYH9 was the gene targets of 126 miRNAs (ex; hsa-mir-181b-3p), APP was the gene targets of 125 miRNAs (ex; hsa-mir-216b-5p), ILF3 was the gene targets of 107 miRNAs (ex; hsa-mir-3157-3p), SKIL was the gene targets 91 of miRNAs (ex; hsa-mir-1294), HSPA8 was the gene targets of 116 of miRNAs (ex; hsa-mir-3661), HSP90AB1 was the gene targets of 103 of miRNAs (ex; hsa-mir-200a-3p), SQSTM1 was the gene targets of 94 of miRNAs (ex; hsa-mir-520d- 5p), HSPA5 was the gene targets of 88 of miRNAs (ex; hsa-mir-573) and GRB2 was the gene targets of 65 of miRNAs (ex; hsa-mir-1291), and are listed in Table 5.

**Fig. 5.**
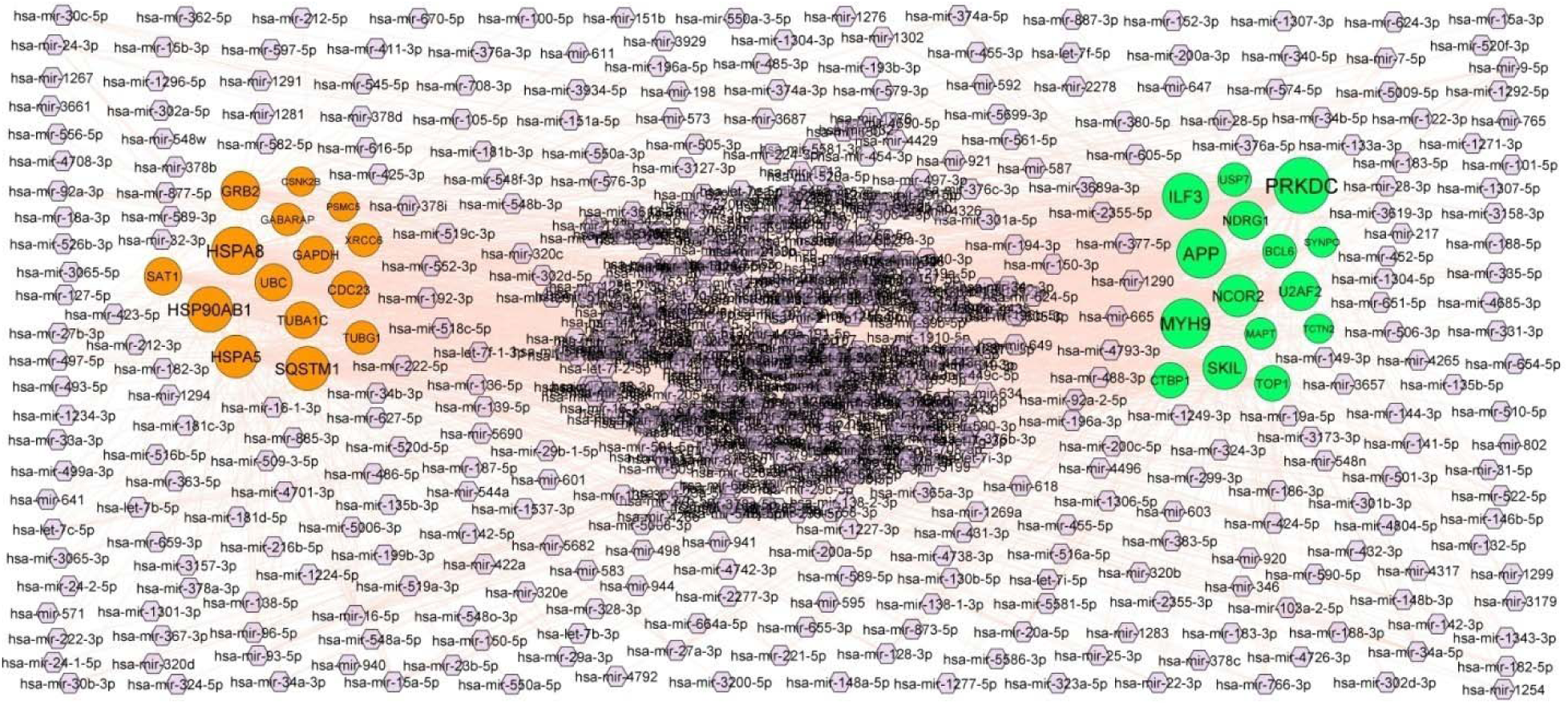
MiRNA - hub gene regulatory network. The chocolate color diamond nodes represent the key miRNAs; up regulated genes are marked in green; down regulated genes are marked in red.

**Table 5.** miRNA - target gene and TF - target gene interaction

| Regulation | Target Genes | Degree | MicroRNA | Regulation | Target Genes | Degree | TF |
| --- | --- | --- | --- | --- | --- | --- | --- |
| Up | PRKDC | 163 | hsa-mir-142-5p | Up | BCL6 | 60 | NOTCH1 |
| Up | MYH9 | 126 | hsa-mir-181b-3p | Up | MYH9 | 53 | PPARD |
| Up | APP | 125 | hsa-mir-216b-5p | Up | NCOR2 | 50 | HIF1A |
| Up | ILF3 | 107 | hsa-mir-3157-3p | Up | APP | 45 | SMARCA4 |
| Up | SKIL | 91 | hsa-mir-1294 | Up | NDRG1 | 44 | SUZ12 |
| Up | NCOR2 | 81 | hsa-mir-4708-3p | Up | TOP1 | 41 | CREM |
| Up | U2AF2 | 65 | hsa-mir-196a-5p | Up | ILF3 | 41 | DACH1 |
| Up | NDRG1 | 60 | hsa-mir-374a-5p | Up | MAPT | 38 | YAP1 |
| Up | TOP1 | 45 | hsa-mir-424-5p | Up | SKIL | 37 | GATA2 |
| Up | CTBP1 | 45 | hsa-mir-944 | Up | PRKDC | 36 | NUCKS1 |
| Up | BCL6 | 35 | hsa-mir-4701-3p | Up | CTBP1 | 35 | ELF1 |
| Up | USP7 | 34 | hsa-mir-93-5p | Up | U2AF2 | 34 | KDM6A |
| Up | MAPT | 23 | hsa-mir-2278 | Up | SYNPO | 28 | SMAD4 |
| Up | SYNPO | 15 | hsa-mir-138-5p | Up | USP7 | 27 | NANOG |
| Up | TCTN2 | 9 | hsa-mir-34a-5p | Up | TCTN2 | 10 | SRF |
| Down | HSPA8 | 116 | hsa-mir-3661 | Down | UBC | 64 | TAF7L |
| Down | HSP90AB1 | 103 | hsa-mir-200a-3p | Down | HSP90AB1 | 49 | RUNX2 |
| Down | SQSTM1 | 94 | hsa-mir-520d-5p | Down | TUBA1C | 47 | MITF |
| Down | HSPA5 | 88 | hsa-mir-573 | Down | HSPA5 | 44 | YAP1 |
| Down | GRB2 | 65 | hsa-mir-1291 | Down | HSPA8 | 39 | E2F1 |
| Down | SAT1 | 56 | hsa-mir-301b-3p | Down | GAPDH | 38 | SOX11 |
| Down | UBC | 55 | hsa-mir-9-5p | Down | GABARAP | 38 | HOXB4 |
| Down | GAPDH | 54 | hsa-mir-5690 | Down | GRB2 | 37 | TFAP2A |
| Down | TUBA1C | 53 | hsa-mir-603 | Down | CSNK2B | 35 | THAP11 |
| Down | CDC23 | 51 | hsa-mir-376a-5p | Down | PSMC5 | 33 | STAT4 |
| Down | TUBG1 | 32 | hsa-mir-182-5p | Down | SQSTM1 | 30 | EP300 |
| Down | XRCC6 | 31 | hsa-mir-618 | Down | CDC23 | 27 | TEAD4 |
| Down | GABARAP | 26 | hsa-mir-34b-3p | Down | SAT1 | 25 | SALL4 |
| Down | PSMC5 | 10 | hsa-mir-452-5p | Down | XRCC6 | 23 | YY1 |
| Down | CSNK2B | 9 | hsa-mir-149-3p | Down | TUBG1 | 22 | SOX2 |

### TF-hub gene regulatory network construction

The hub genes of the DEGs in T2DM were performed by online databases NetworkAnalyst. Based on the TFs, a TF -hub gene regulatory network was constructed with 477 nodes (TF: 192 and hub gene: 285) and 8507 interaction pairs (Fig. 6). BCL6 was the gene targets of 60 TFs (ex; NOTCH1), MYH9 was the gene targets of 53 TFs (ex; PPARD), NCOR2 was the gene targets of 50 TFs (ex; HIF1A), APP was the gene targets of 45 TFs (ex; SMARCA4), NDRG1 was the gene targets of 44 TFs (ex; SUZ12), UBC was the gene targets of 64 TFs (ex; TAF7L), HSP90AB1 was the gene targets of 49 TFs (ex; RUNX2), TUBA1C was the gene targets of 47 TFs (ex; MITF), HSPA5 was the gene targets of 44 TFs (ex; YAP1) and HSPA8 was the gene targets of 39 TFs (ex; E2F1), and are listed in Table 5.

**Fig. 6.**
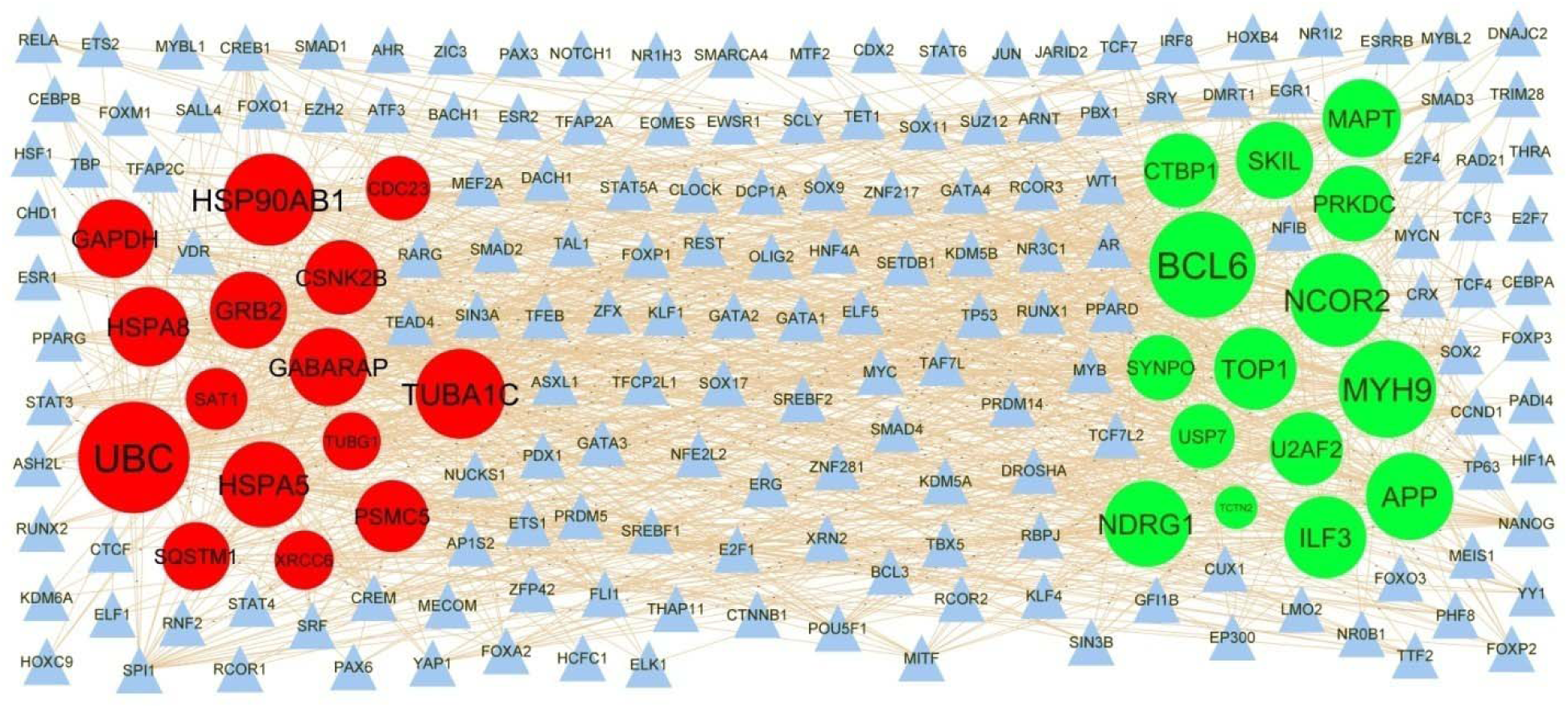
TF - hub gene regulatory network. The blue color triangle nodes represent the key TFs; up regulated genes are marked in green; down regulated genes are marked in red.

### Validation of hub genes by receiver operating characteristic curve (ROC) analysis

Validated by ROC curves, we found that 10 hub genes had high sensitivity and specificity, including APP (AUC□=□0.853), MYH9 (AUC□=□0.852), TCTN2 (AUC□=□0.881), USP7 (AUC□=□0.862), SYNPO (AUC□=□0.893), GRB2 (AUC□=□0.850), HSP90AB1 (AUC□=□0.870), UBC (AUC□=□0.865), HSPA5 (AUC□=□0.902) and SQSTM1 (AUC□=□0.875) (Fig. 7). The hub genes might be biomarkers of T2DM and have positive implications for early medical intervention of the disease.

**Fig. 7.**
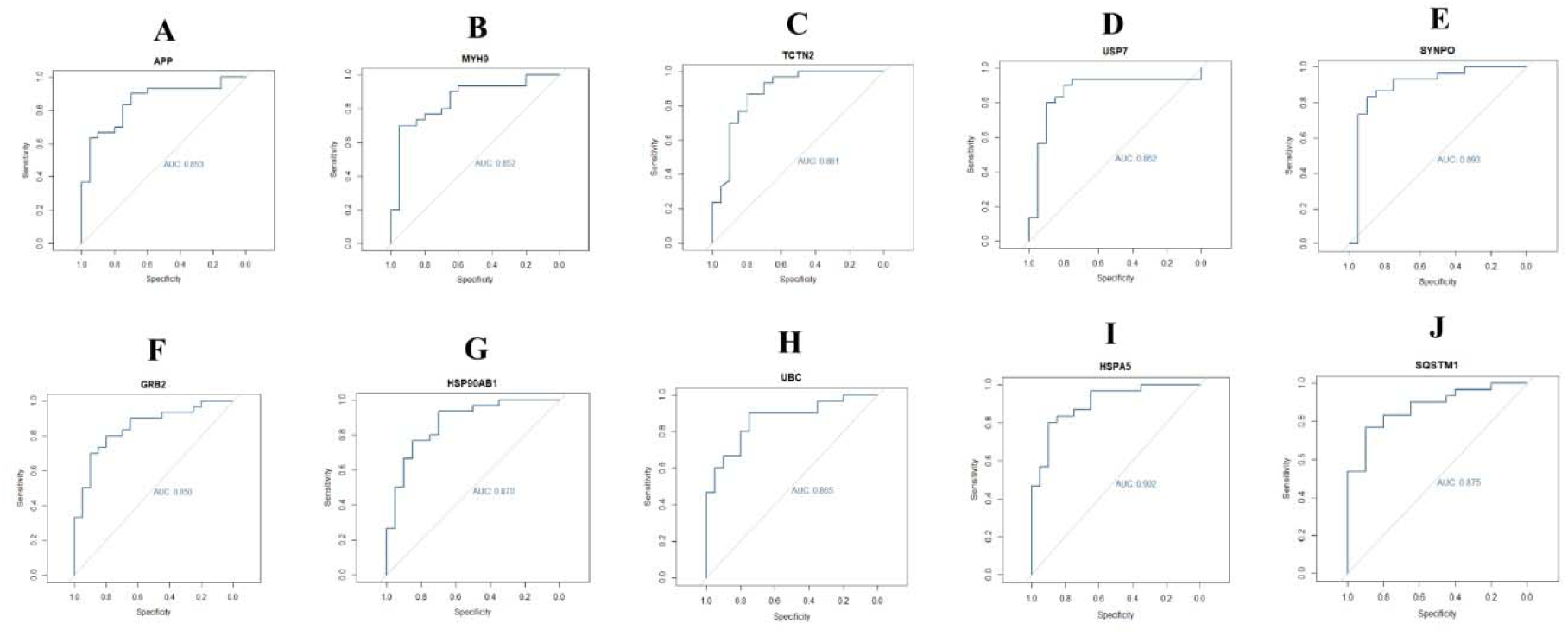
ROC curve validated the sensitivity, specificity of hub genes as a predictive biomarker for T2DM. A) APP B) MYH9 C) TCTN2 D) USP7 E) SYNPO F) GRB2 G) HSP90AB1 H) UBC I) HSPA5 J) SQSTM1

## Discussion

Although there are various investigations on T2DM have been conducted, the mortality of T2DM is still high. This might be due to the lack of valid biomarkers for detection of early stage T2DM and of valid treatment for T2DM. Therefore, molecular mechanisms of T2DM are necessary for scientists to find the treat and diagnosis method of T2DM. Because of the fast advancement of RNA sequencing technology, it is more convenient to find out the genetic modification of development of diseases. RNA sequencing facilitates us to examine the gene, the genetic modification in T2DM, which had been proved to be a better approach to find novel biomarkers in other metabolic diseases.

In the present investigation, we observed whether there were more beneficial genes which could be better biomarkers for the diagnosis, prognosis and therapeutic for T2DM. In order to find out the significant gene of T2DM, we analyzed the T2DM expression profiling by high throughput sequencing of GSE81608 in limma, where a total number of 927 DEGs were obtained between T2DM and normal control, comprising was 461 up regulated and 466 down regulated genes. CTBP1 [35] and TRNC [36] are involved in the pathogenesis of T2DM. Previous studies have demonstrated that SST (somatostatin) serve an essential role in obesity [37], but this gene might be novel target for T2DM.

Then, databases including GO and REACTOME were selected to do gene enrichment analysis. Metabolism of proteins [38], metabolism [39], the citric acid (TCA) cycle and respiratory electron transport [40], gluconeogenesis [41], immune system [42], heterocyclic compound binding [43], protein binding [44], establishment of localization [45], cellular metabolic process [46], cytoplasm [47] and catalytic activity [48] were the GO terms and signaling pathways responsible for the advancement of T2DM. A previous study showed that IGFBP2 [49], APOH (apolipoprotein H) [50], ANXA2 [51], BAX (BCL2 associated X, apoptosis regulator) [52], PCSK1N [53], PDK4 [54], CPE (carboxypeptidase E) [55], OCLN (occludin) [56], CD44 [57], NDN (necdin, MAGE family member) [58], MLXIPL (MLX interacting protein like) [59], CD36 [60], SREBF1 [61], NR4A1 [62], PCSK2 [63], CHGB (chromogranin B) [64], PDK3 [65], PDCD4 [66], EIF5A [67], NRP1 [68], ABCA1 [69], DNMT1 [70], MYH9 [71], HMGB1 [72], B4GALT5 [73], B2M [74], MAP3K12 [75], KSR2 [76], NPY (neuropeptide Y) [77], CHGA (chromogranin A) [78], CD47 [79], DLK1 [80], PDK4 [81], CPE (carboxypeptidase E) [82], OCLN (occludin) [83], CXXC4 [84], PEMT (phosphatidylethanolamine N-methyltransferase) [85], FADS2 [86], RREB1 [87], HNRNPAB (heterogeneous nuclear ribonucleoprotein A/B) [88], CPT1A [89], ALDH1B1 [90], ESRRA (estrogen related receptor alpha) [91], NISCH (nischarin) [92], SSTR3 [93], ND1 [94], NCOR2 [95], RBP4 [96], GSTP1 [97], CYB5A [98], G6PC2 [99], DNAJC15 [100], TMED6 [101], PSMD6 [102], CLU (clusterin) [103], TTR (transthyretin) [104], TXN (thioredoxin) [105], LAMTOR1 [106], GLUL (glutamate-ammonia ligase) [107], NEU1 [108], HSPA8 [109], AP3S2 [110], COX4I1 [111], MT2A [112] MTCH2 [113], ESD (esterase D) [114], UBE2L6 [115], SCD (stearoyl-CoA desaturase) [116], MGST3 [117], NQO1 [118] NSMCE2 [119] and PRSS1 [120] played an important role in T2DM. Quintela et al [121], Yuan et al [122], Cacace et al [123], Hao et al [124], Beckelman et al [125], Liu et al [126], Sekiguchi et al [127], Castillon et al [128], O’Donnell-Luria et al [129], Coupland et al [130], Koufaris et al [131], Qvist et al [132], Richter et al [133], Torres et al [134], Jeong et al [135], Bermejo-Bescós et al [136], Ramon-Duaso et al [137], Guilarte, [138], Mukaetova-Ladinska et al [139], Fazeli et al [140], Butler et al [141], Nackenoff et al [142], Konyukh et al [143], Hu et al [144], Kaur et al [145], Nakamura et al [146], Liu et al [147], Obara et al [148], Herrmann et al [149], Ozgen et al [150], Masciullo et al [151], Perrone et al [152], Su et al [153], Zhao et al [154], Iqbal et al [155], Gal et al [156], Wang et al [157], Stefanović et al [158], Zahola et al [159], Bik-Multanowski et al [160], Mata et al [161], Li et al [162], Payton et al [163] and Chai et al [164] indicated that UBA6, TIA1, DPP6, USP7, EEF2, ITM2B, DPH1, PAK3, KMT2E, MAPT (microtubule associated protein tau), HCFC1, BRD1, TAOK2, PHF1, STMN2, APP (amyloid beta precursor protein), MBNL2, APLP1, MAP2, SRRM2 CST3, SRRM2, CST3, PLD3, SEZ6L2, DOC2A, PI4KA, GNAO1, TRA2A, MIDN (midnolin), HOOK3, MCPH1, SACS (sacsin molecular chaperone), TUBA4A, ASAH1, ATP6V1B2, SVBP (small vasohibin binding protein), AIFM1, UBC (ubiquitin C), IFI30, SCGN (secretagogin, EF-hand calcium binding protein), MTRNR2L12, GBA (glucosylceramidase beta), TXN2, NQO2 and PPIL1 were involved in the development and progression of cognitive impairment, but these genes might be novel target for T2DM. RPS3A [165], PGAM5 [166], RPL7 [167], TLK1 [168], DDR1 [169], ILF3 [170], TNRC6A [171], GGCX (gamma-glutamyl carboxylase) [172], S100A6 [173], LSAMP (limbic system associated membrane protein) [174], KCNA5 [175], LUC7L3 [176], ATAD3C [177], SRSF3 [178], MCU (mitochondrial calcium uniporter) [179], ATP2A2 [180], GAA (glucosidase alpha, acid) [181], MAGI1 [182], WIPF2 [183], VAMP8 [184], UCHL1 [185], CLIC1 [186], PSMB5 [187], GRB2 [188], ZMPSTE24 [189], COX6B1 [190], SQSTM1 [191], COTL1 [192], CD63 [193], NDUFB7 [194], BEX1 [195] and MTRNR2L8 [196] plays a major role in mediating cardiovascular diseases progression, but these genes might be novel target for T2DM. HLA-A [197], VEGFA (vascular endothelial growth factor A) [198], RPS26 [199], BMP6 [200], HLA-B [201], IER3IP1 [202], MT1E [203], ACADM (acyl-CoA dehydrogenase medium chain) [204] and GAPDH (glyceraldehyde-3-phosphate dehydrogenase) [205] are associated with progression of type 1 diabetes mellitus, but these genes might be novel target for T2DM. PEMT (phosphatidylethanolamine N-methyltransferase) [206], INSM1 [207], BCL6 [208], RUNX1T1 [209], PGRMC2 [210], ARID1B [211], CITED2 [212], KLF13 [213], PPT1 [214], ARRDC3 [215], HSPA5 [216], MDH2 [217] and COA3 [218] have been previously reported to be a key biomarkers for the early detection of obesity, but these genes might be novel target for T2DM. A previous study demonstrated that IGFBP5 [219], PRDX6 [220], PKM (pyruvate kinase M1/2) [221], PRDX1 [222] and USP22 [223] were more highly expressed in diabetic nephropathy, but these genes might be novel target for T2DM. Durgin et al [224], Zhang et al [225], Hamada et al [226], Gong et al [227], Li et al [228], Lin et al [229] and Schweigert et al [230] suggested that CYB5R3, CACNA1A, GLCCI1, CAP1, HSP90AB1, BLVRA (biliverdinreductase A) and CRIP1 were involved in the progression of hypertension, but these genes might be novel target for T2DM.

By PPI network and module analysis, we identified the hub genes that might affect the origin or advancement of T2DM. TCTN2, SYNPO (synaptopodin), PSMD12, PSMC4, TUBA1C, PSMC5, PSMD7 and RAD23A were novel biomarkers for the progression of T2DM.

In addition, miRNA-hub gene regulatory network construction and TF-hub gene regulatory network were constructed. In addition, miRNA-mRNA networks were constructed. The roles of hub genes, miRNA and TF in the pathogenesis of T2DM are discussed. Hsa-mir-142-5p [231], hsa-mir-1291 [232], NOTCH1 [233], PPARD (peroxisome proliferator-activated receptor delta) [234], HIF1A [235], RUNX2 [236] and E2F1 [237] levels are correlated with disease severity in patients with T2DM. Previous studies have demonstrated that hsa-mir-216b-5p [238] and hsa-mir-200a-3p [239] appears to be expressed in type 1 diabetes, but these genes might be novel target for T2DM. Hsa-mir-1294 [240], SUZ12 [241] and YAP1 [242] were responsible for progression of cognitive impairment, but these genes might be novel target for T2DM. Hsa-mir-573 [243] and SMARCA4 [244] were linked with progression of hypertension, but these genes might be novel target for T2DM. PRKDC (protein kinase, DNA-activated, catalytic subunit), SKIL (SKI like proto-oncogene), NDRG1, hsa-mir-181b-3p, hsa-mir-3157-3p, hsa-mir-3661, hsa-mir-520d-5p, TAF7L and MITF (Microphthalmia-associated transcription factor) were novel biomarkers for the progression of T2DM.

In conclusion, the present study identified ten hub genes (APP, MYH9, TCTN2, USP7, SYNPO, GRB2, HSP90AB1, UBC, HSPA5 and SQSTM1) with crucial role in progression of T2DM; our results suggested these genes could add a new dimension to our understanding of the T2DM and might be served as potential biomarkers that will be assisting endocrinologist in developing novel therapeutic strategies for T2DM patients. However, there are some limitations in this study. Further larger clinical sample size and in-depth clinical experiment are needed to clarify the clear mechanism and warrant the prognostic value of these DEGs in T2DM.

## Acknowledgement

I thank Yurong Xin, Regeneron Pharmaceuticals, Inc., Tarrytown, New York, USA, very much, the author who deposited their profiling by high throughput sequencing dataset GSE81608, into the public GEO database.

## Conflict of interest

The authors declare that they have no conflict of interest.

## Ethical approval

This article does not contain any studies with human participants or animals performed by any of the authors.

## Informed consent

No informed consent because this study does not contain human or animals participants.

## Availability of data and materials

The datasets supporting the conclusions of this article are available in the GEO (Gene Expression Omnibus) (https://www.ncbi.nlm.nih.gov/geo/) repository. [(GSE81608) (https://www.ncbi.nlm.nih.gov/geo/query/acc.cgi?acc=GSE81608)]

## Consent for publication

Not applicable.

## Competing interests

The authors declare that they have no competing interests.

## Author Contributions

V. A - Methodology and validation

V. R - Formal analysis and validation

B. V - Writing original draft, and review and editing

C. V - Software and investigation

S. K - Supervision and resources

## Notes

### Competing Interest Statement

The authors have declared no competing interest.

